# Monomethylation of Lysine 27 at Histone 3 Confers Lifelong Susceptibility to Stress

**DOI:** 10.1101/2023.05.08.539829

**Authors:** Angélica Torres-Berrío, Molly Estill, Aarthi Ramakrishnan, Hope Kronman, Vishwendra Patel, Angélica Minier-Toribio, Orna Issler, Caleb J. Browne, Eric M. Parise, Yentl van der Zee, Deena Walker, Freddyson J. Martínez-Rivera, Casey K. Lardner, Romain Durand-de Cuttoli, Scott J. Russo, Li Shen, Simone Sidoli, Eric J. Nestler

## Abstract

Histone post-translational modifications are critical for mediating persistent alterations in gene expression. By combining unbiased proteomics profiling, and genome-wide approaches, we uncovered a role for mono-methylation of lysine 27 at histone H3 (H3K27me1) in the enduring effects of stress. Specifically, mice exposed to early life stress (ELS) or to chronic social defeat stress (CSDS) in adulthood displayed increased enrichment of H3K27me1, and transient decreases in H3K27me2, in the nucleus accumbens (NAc), a key brain-reward region. Stress induction of H3K27me1 was mediated by the VEFS domain of SUZ12, a core subunit of the polycomb repressive complex-2, which is induced by chronic stress and controls H3K27 methylation patterns. Overexpression of the VEFS domain led to social, emotional, and cognitive abnormalities, and altered excitability of NAc D1 mediums spiny neurons. Together, we describe a novel function of H3K27me1 in brain and demonstrate its role as a “chromatin scar” that mediates lifelong stress susceptibility.

## INTRODUCTION

A life history of stress is the strongest known risk factor for depression and anxiety, which are among the world’s leading causes of disability^1, 2^. Indeed, stress is known to induce persistent transcriptional and cellular signatures in regions involved in reward and mood regulation^3–5^. These effects can be observed long after stress exposure has ceased and are highly sensitive to windows of brain neuroplasticity, resulting in several behavioral abnormalities that range from heightened stress reactivity and social avoidance to cognitive dysfunction and anhedonia^2, 6–8^. Despite current efforts to uncover the mechanisms that underlie the behavioral, cellular and transcriptional phenotypes induced by stress, there is still a lack of information about the molecular players that mediate these long-lasting abnormalities.

The nucleus accumbens (NAc), a core component of the brain’s reward circuitry, serves as a hub region that integrates information related to reward, motivation and complex cognitive function^9, 10^. By receiving inputs from multiple brain areas, the NAc not only fine-tunes behavioral outputs towards rewarding and away from aversive stimuli^11^, but also determines appropriate responses to stressful events^3, 12, 13^. These responses are shaped by two subtypes of medium spiny projection neurons (MSNs): D1 and D2 dopamine receptor-expressing cells^14–16^, which are segregated, both transcriptionally and electrophysiologically, and are critical for the development of susceptible vs resilient stress-related phenotypes^14–16^.

Using genome-wide RNA-sequencing (RNA-seq) in postmortem brain tissue of individuals with depression and mouse models of chronic stress, our group has identified broad transcriptional changes in the NAc and its innervating regions^4, 5, 17^. Interestingly, these changes in gene expression can be further exacerbated by a previous history of early life stress (ELS)^6, 8^, suggesting latent mechanisms that could promote or repress future transcriptional states. Converging evidence supports the hypothesis that histone post-translational modifications are critical for stress-induced changes in gene expression^2, 6, 18–20^. Such regulation occurs through the addition or removal of methyl, acetyl or other groups at specific histone tail residues, which together orchestrate chromatin remodeling and ultimately dictate the availability of DNA for lasting transcription^21^. However, while certain histone modifications have been implicated in stress-related disorders, their influence is restricted to a relatively small number of genes and may not be recapitulated across different stress models^19, 20^. In this context, the use of unbiased proteomic approaches has become a powerful tool to unravel those histone modifications that are most strongly affected in NAc in response to environmental events, such as stress, and that may influence the broad transcriptional states in this brain region^6^.

Here, we combined mass spectrometry profiling, genome-wide approaches, and mouse models of stress to demonstrate the role of mono-methylation of lysine 27 at histone H3 (H3K27me1) acting in the NAc in conferring persistent vulnerability to stress. We report increased levels of H3K27me1 in the NAc of susceptible, but not resilient, mice after chronic social defeat stress (CSDS). Moreover, this effect is recapitulated in adolescent and adult mice that were previously exposed to ELS. Stress-induced enrichment of H3K27me1 in the NAc is associated with induction of SUZ12, a core subunit of the polycomb repressive complex-2 (PRC2), which determines H3K27 methylation patterns. Indeed, overexpression of the VEFS domain of SUZ12 in D1-MSNs mimics the selective induction of H3K27me1 and leads to persistent behavioral and electrophysiological signatures of stress susceptibility. This study thereby establishes a novel function of H3K27me1 in brain, and demonstrates its role as an important “chromatin scar” that mediates lifelong stress susceptibility.

## RESULTS

### Dynamic methylation of H3K27 in response to social stress

To identify global changes in histone modifications associated with stress susceptibility, we used mass spectrometry, an unbiased proteomic approach that allows for the quantification of the relative abundance of hundreds of histone marks. We subjected adult male mice to CSDS^22^, a validated model for the study of depression-like behaviors that separate mice into susceptible and resilient populations based on a social interaction test, which is highly predictive of numerous other behavioral abnormalities^23^. Bulk NAc tissue from control, susceptible and resilient male mice was collected 24 hours after the social interaction test and processed for mass spectrometry (Figure 1A-B).

**Figure 1.**
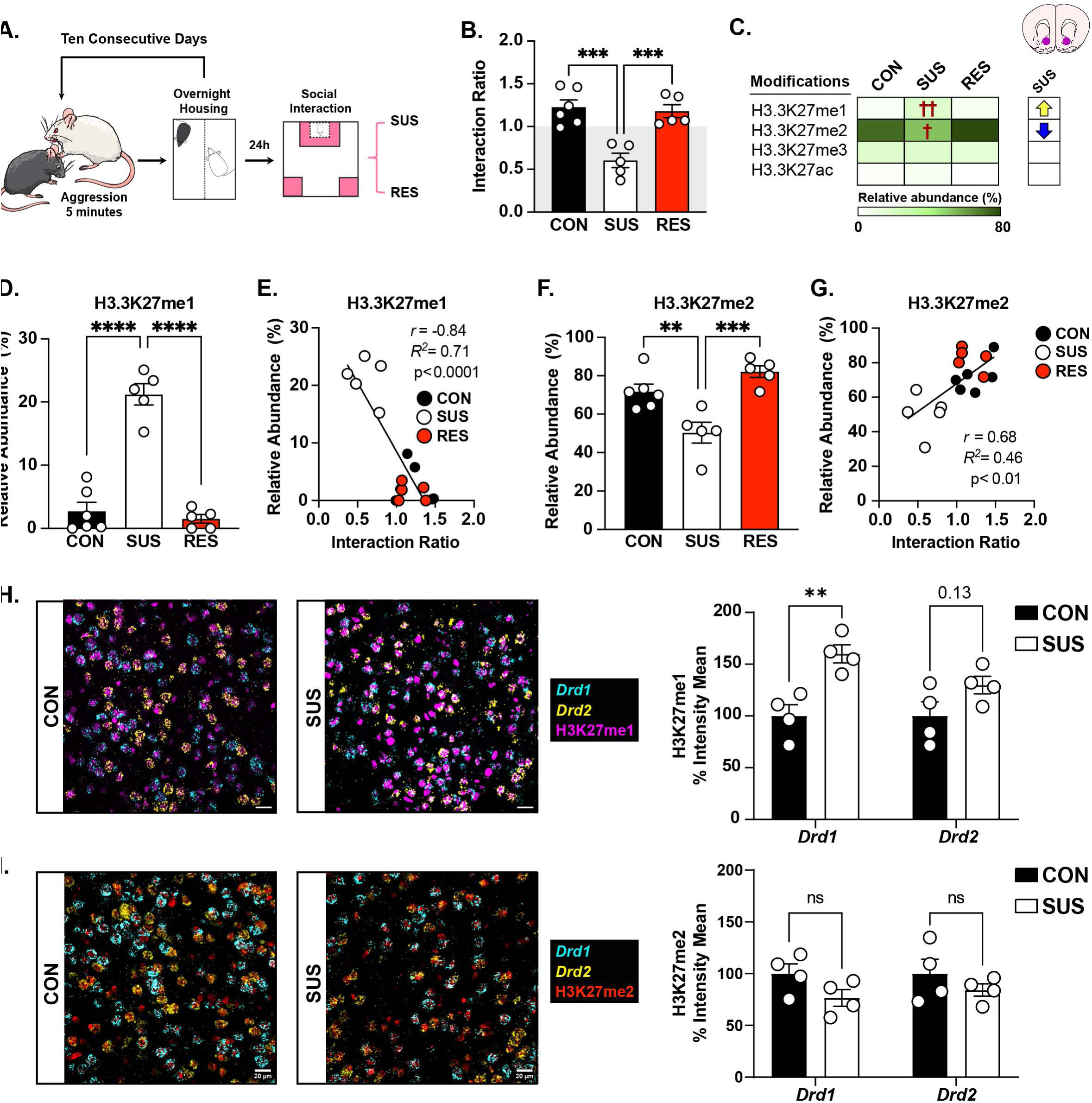
Dynamic regulation of H3K27 methylation in NAc after CSDS. **(A)** Schematic illustration and timeline of the CSDS experiment. **(B)** Social interaction ratio: One-way ANOVA: F_(2,13)_=17.32, p=0.0002. Tukey’s test: susceptible (SUS, n=5) different from control (CON, n=6) and resilient (RES, n=5) mice, ****p<0.001. **(C)** Heatmap display of the relative abundance of H3K27 modifications in CON, SUS and RES mice. The data was obtained by mass spectrometry. Arrows on the right indicate significant p-values for one-way ANOVA of each mark (yellow, increase: ^††^p<0.0001; blue, decrease: ^†^p<0.001). **(D)** H3.3K27me1 relative abundance: One-way ANOVA: F_(2,13)_=65.21; p<0.0001. Tukey’s test: Increased H3.3K27me1 in SUS mice compared to CON and RES,****p<0.00001. **(E)** Negative correlation between H3.3K27me1 abundance and social interaction ratio. **(F)** H3.3K27me2 relative abundance: One-way ANOVA: F_(2,13)_= 14.15, p=0.0005. Tukey’s test: Decreased H3.3K27me2 in SUS mice compared to CON (**p<0.001) and RES (***p<0.0001) mice. **(G)** Positive correlation between H3.3K27me2 abundance and social interaction ratio. **(H)** Expression of H3K27me1 in NAc MSNs (n=3-4 bilateral sections; 4 mice/group). Left: Representative coronal section of CON and SUS mice showing H3K27me1 expression (magenta) in *Drd1-* (cyan) and *Drd2-* (yellow) MSNs. Right: Two-way ANOVA: group effect: F_(1,12)_=17.87; p=0.0012; no significant cell-type effect: F_(1,12)_=2.04; p=0.17, or interaction: F_(1,12)_= 2.02; p=0.18. Sidak’s comparison for group effect: SUS different from CON in *Drd1-MSNs*, **p<0.0036**. (I)** Expression of H3K27me2 in NAc MSNs. Left: Representative coronal section of CON and SUS mice showing H3K27me2 expression (red) in *Drd1-* and *Drd2-*MSNs. Right: Two-way ANOVA: no significant effect of group: F_(1,12)_=4.07; p=0.07; cell-type: F_(1,12)_=0.156; p=0.69, or interaction: F_(1,12)_=0.15; p=0.705. Scale bar: 20µm.

Analysis of the relative abundance of single peptides revealed significant changes in a modest number of histone modifications in the NAc of susceptible and resilient mice, including mono-, di- or trimethylation (me1, me2, or me3) at K9, K27, and K36 of H3 (Figure S1A). Strikingly, we found that susceptible mice displayed the greatest alterations in methylation dynamics of K27 of the histone variant H3.3 (Figure 1C): there was a significant increase in the abundance of H3.3K27me1 (Figure 1D), and a corresponding decrease of H3.3K27me2 (Figure 1F). Furthermore, we identified that the levels of both histone marks were highly correlated, albeit in opposite directions, with social interaction (Figure 1E-G). By contrast, H3.3K27me3, associated with “repressive” chromatin, and H3.3K27ac, a mark for active enhancers, were unchanged in susceptible or resilient groups as compared to control mice (Figure 1C). Regulation of H3.3K27me1 and -me2 was specific to males, as we did not find differences in female mice after CSDS (Figure S1B-E). Therefore, we focused on male mice moving forward.

H3.3 is the dominant form of H3 present in adult brain neurons, where it represents up to 90% of total histone H3^24–26^. The H3.3 variant contains a serine (S) for alanine (A) substitution at position 31 of the N-terminus (Figure S1F)^26^. The methylation of K27 at H3, and all of its variants, is catalyzed by PRC2^27–29^. Given that currently available antibodies recognize K27me1 and K27me2 in H3 and H3.3 (Figure S1F), from here onward we refer to these histone marks as H3K27me1 and H3K27me2.

We found that H3K27me1 and H3K27me2 are primarily expressed by D1 or D2 MSNs in the NAc compared to other cell types in this region (Figure S1G-H). We next assessed whether CSDS-induced changes in these histone marks are cell-specific. To this end, we combined RNAscope with immunofluorescence to analyze H3K27me1 and H3K27me2 levels in D1 or D2 MSNs of control and susceptible mice. We found that susceptible mice exhibited a significant increase in H3K27me1 levels, an effect that was driven mainly by D1-MSNs in the NAc (Figure 1H). In contrast, we observed a non-significant trend toward decreased H3K27me2 levels in both D1 and D2 MSNs (Figure 1I). These results confirm the opposite expression of the two histone marks in the NAc of CSDS-susceptible mice, and indicate that H3K27me1 upregulation occurs primarily in D1-MSNs.

### H3K27me1 is deposited across intragenic regions of genes associated with neuronal excitability

To determine the distribution of H3K27me1 and H3K27me2 across the genome and identify putative target genes, we used Cleavage Under Targets & Release Using Nuclease followed by DNA-sequencing (CUT&RUN-seq)^30^, using validated antibodies for H3K27me1 and H3K27me2 (Figure S2A-D). We collected bulk NAc tissue punches from control, susceptible and resilient mice 24 hours after the social interaction test and isolated nuclei for CUT&RUN-seq (Figure 2A). We first identified genomic regions to which H3K27me1 and H3K27me2 bind under control conditions by annotating the peaks based on their location in the mouse reference genome (mm10). We confirmed previous observations from non-nervous tissues^31, 32^ that both histone marks are deposited across non-overlapping sites within the mouse genome. Indeed, the vast majority of H3K27me1 accumulated within gene bodies, followed by promoters, whereas H3K27me2 was predominantly enriched across intergenic regions (Figure S2E-F).

**Figure 2.**
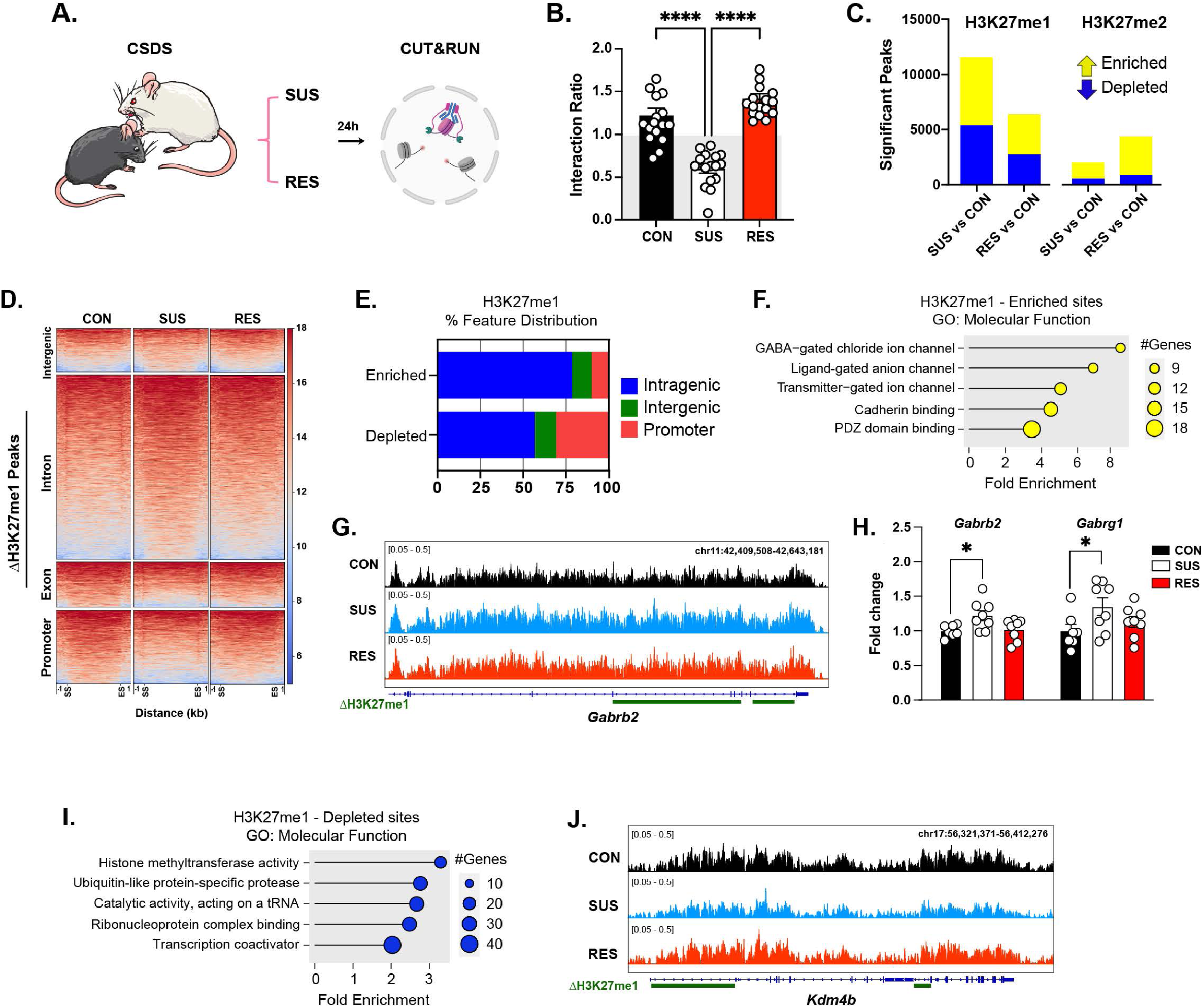
CUT&RUN analysis of the genome-wide deposition of H3K27me1 and H3K27me2 in NAc under control and CSDS conditions. **(A)** Schematic and timeline of CUT&RUN-seq experiment (n=4 samples, each consisting of pooled NAc tissue from 4 mice from the same group). Validation of the antibodies in Figure S2. **(B)** Social interaction ratio of CON, SUS and RES mice (n=16/group). One-way ANOVA: F_(2,45)_=39.49; p<0.0001. Tukey’s test: SUS different from CON and RES mice, ****p<0.0001. **(C)** Number of differential peaks enriched (yellow) of depleted (blue) in H3K27me1 and H3K27me2 in the SUS vs CON, and RES vs CON comparisons. **(D)** Coverage heatmap for H3K27me1 enrichment within a region spanning ± 1 kb around the start site (SS) and end site (ES) of the differential peaks in the SUS vs CON comparison. The gradient blue-to-red color indicates low-to-high counts within intragenic, promoter, exon and intergenic regions. Coverage heatmap for H3K27me2 is shown in Figure S2H. **(E)** Genomic distribution of H3K27me1 differential peaks in NAc in the SUS vs CON comparison. The data show that stress-induced enrichment of H3K27me1 dominates across intragenic regions, while depletion is observed within intragenic and promoter regions. **(F)**. Top 5 GO molecular function terms associated with H3K27me1 enrichment, ranked according to their log-10(FRD), followed by fold enrichment. Additional terms are presented in Table S1 **(G)** Representative Integrative Genomics Viewer (IGV) browser tracks of H3K27me1 enrichment across the Gamma-aminobutyric acid type A receptor subunit beta2, *Gabrb2*, in CON, SUS and RES mice. ΔH3K27me1 sites span across introns 4, 5, 6 and 9. **(H)** Quantitative PCR showing increased of *Gabrb2* and *Gabrg1* gene expression in SUS mice. *Gabrb2*: One-way ANOVA: F_(2,20)_=4.46, p<0.05. Tukey’s test: SUS different from CON mice, *p<0.05. *Gabrg1*: One-way ANOVA: One-way ANOVA: F_(2,20)_= 4.46, p<0.05. Tukey’s test: SUS different from CON mice, *p<0.05. **(I)** Top 5 GO molecular function terms associated with H3K27me1 depletion. Other terms are presented in Table S2. **(J)** Representative track of H3K27me1 depletion across the lysine demethylase 4B *(Kdm4b)* gene in CON, SUS and RES mice. ΔH3K27me1 sites span across introns 1 and 9.

To better understand the contribution of H3K27me1 and H3K27me2 to the actions of CSDS, we analyzed differentially enriched sites of each mark within susceptible or resilient groups relative to non-defeated controls. We found a greater number of H3K27me1 differential peaks in the susceptible vs control comparison, with several thousands of enriched (5254) and depleted (5180) sites, than in the resilient vs control comparison (Figure 2C). By contrast, the number of enriched sites for H3K27me2 was much lower in the susceptible vs control comparison, with only 1440 upregulated peaks, and a modestly larger number in the resilient vs control comparison (3706 upregulated peaks) (Figure 2C). We then analyzed the genomic distribution of differentially-expressed H3K27me1 and H3K27me2 peaks in the susceptible vs control comparison. We found that changes in H3K27me1 enrichment generally mirrored the baseline distribution of the mark, with ∼90% of regulated sites being within intragenic or promoter regions, although far more depletion than enrichment occurred at promoters (Figure 2D-E), which suggests that some H3K27me1-enriched sites are more accessible than others to mechanisms that regulate K27 methylation^32^. Changes in H3K27me2, on the other hand, were mainly restricted to intergenic regions (∼87%), with very little regulation seen within gene bodies or promoters (Figure S2G-H). We focused on H3K27me1 for further downstream analyses.

Given the much larger number of differential peaks for H3K27me1 in the susceptible vs control comparison, we performed gene ontology (GO) analysis to examine known molecular functions affected under these conditions. Increased H3K27me1 deposition was associated with terms such as *GABA function*, *ion channel activity* and overall neuronal excitability (Figure 2F, Table S1). This finding was further supported by the GO cellular component, reporting “*GABA-ergic synapse*” as the most significantly enriched term (Table S1). Some of the genes that encode for GABA receptor subunits are clustered within neighboring loci and exhibit high H3K27me1 deposition, with complete depletion of H3K27me2 (Figure S3F), suggesting that stress-induced enrichment of H3K27me1 may affect the transcriptional potential of genes involved in GABA-related neurotransmission. Indeed, expression of the beta 2 (*Gabrb2)* and gamma 1 *(Gabrg1)* subunits of the GABA_A_ receptor was significantly increased in the NAc of susceptible mice after CSDS (Figure 2G-H).

Finally, we conducted a GO analysis on the gene sites displaying <u>reduced</u> deposition of H3K27me1 after CSDS, and identified molecular functions associated with *histone methyltransferase activity*, and overall transcription and posttranscriptional regulation (Figure 2I, Table S2). We observed that several genes previously implicated in depression^33, 34^, such as lysine-specific histone demethylase-4B (*Kdm4b,* Figure 2J), or the microRNA regulator *Dicer1*, displayed less H3K27me1 accumulation in susceptible mice (Table S2). Overall, these findings confirm our proteomics dataset and establish patterns of H3K27me1 regulation in the NAc of susceptible mice. Furthermore, they indicate that different molecular functions can be affected by the enrichment vs depletion of H3K27me1 in this brain region.

### VEFS domain of SUZ12 induces H3K27me1 in D1-MSNs and confers susceptibility to stress

Methylation dynamics of H3K27, including its H3.3 variant, are catalyzed by PRC2^27–29^, a complex that involves the coordinated action of three core proteins: enhancer of Zeste 2 (EZH2), suppressor of Zeste 12 (SUZ12), and embryonic ectoderm development (EED)^27, 35, 36^, and that displays stronger preference for H3K27me1 and unmethylated substrates *in vitro*^37^. To assess the potential role of these core PRC2 subunits in stress susceptibility, we measured their expression levels in the NAc of mice exposed to CSDS. We found elevated levels of SUZ12 in susceptible mice compared to their control counterparts (Figure 3A), with no differences in EZH2 or EED observed (Figure 3B-C).

**Figure 3.**
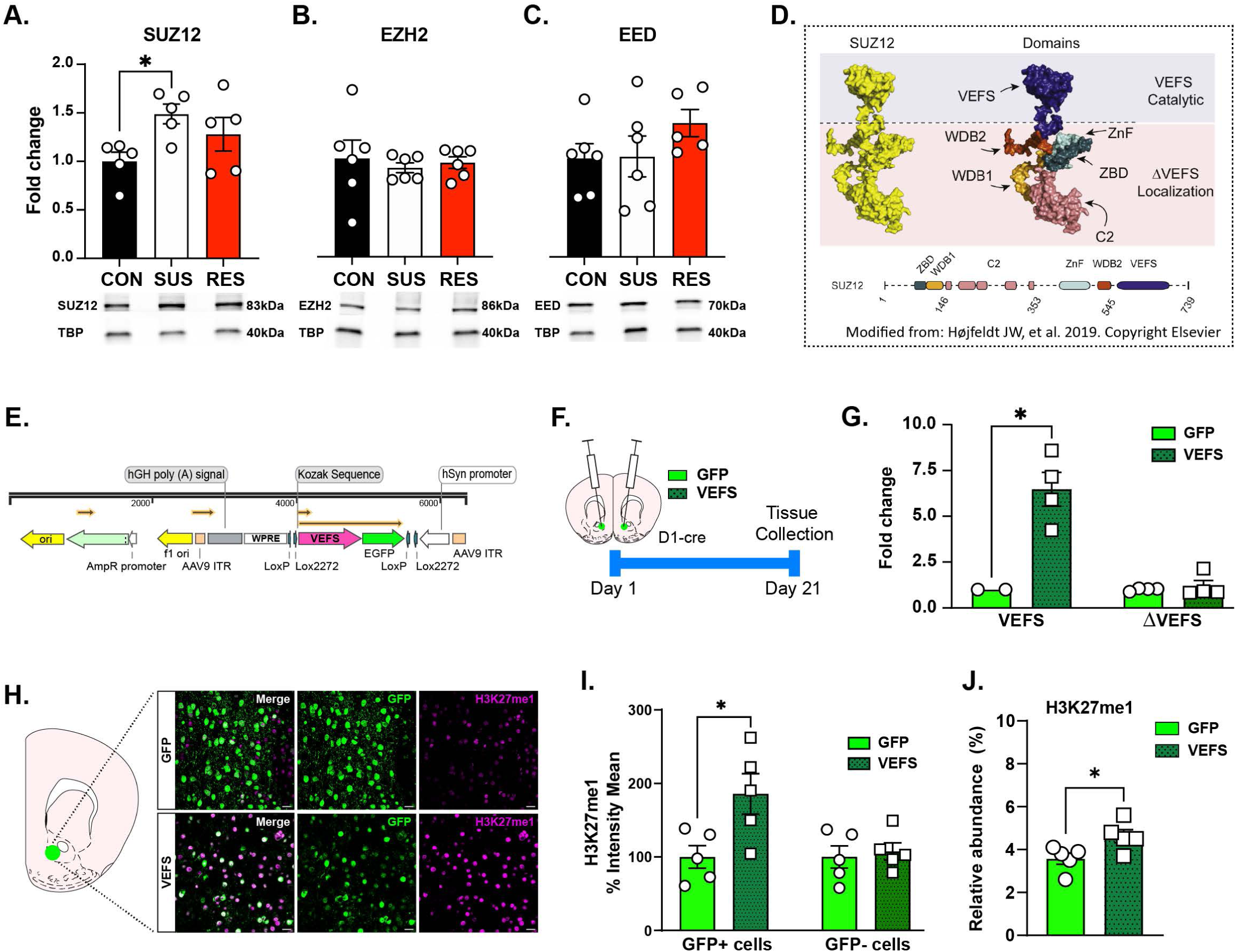
VEFS domain of SUZ12 induces H3K27me1 in NAc D1-MSNs. **(A)** Increased SUZ12 expression in SUS mice compared to CON. One-Way ANOVA: F_(2,12)_=3.62; p<0.05. Tukey’s test: SUS different from CON mice, *p<0.05. **(B)** No significant differences in EZH2 protein levels: One-Way ANOVA: F_(2,15)_=0.18; p=0.83. **(C)** No significant differences in EED protein levels: One-Way ANOVA: F_(2,14)_=1.28; p=0.3. Tissue samples of CON, SUS and RES mice were collected from the same CSDS experiments (n=5-6/group). **(D)** Left: Schematic of the SUZ12 protein (in yellow). Right: Structure of SUZ12, colored to highlight its catalytic (VEFS) and localization (ΔVEFS) domains. The VEFS catalytic domain (in blue) determines the interaction between SUZ12 and EZH2. Modified with permission from Højfeldt, et al. (2019)^50^. Copyright Elsevier. **(E)** Schematic of the Cre-dependent VEFS overexpression AAV construct. **(F)** Timeline of viral infection experiment. **(G)** Quantitative PCR amplification of primers designs for the VEFS or ΔVEFS domains in the NAc of mice injected with either AAV-GFP or AAV-VEFS viruses under basal conditions (n=4/group). VEFS: t_(4)_=3.95, p<0.05; ΔVEFS: t_(6)_=0.57, p=0.58. **(H)** Representative coronal section of a stress-naive mouse showing H3K27me1 expression in AAV-GFP-injected (top) or AAV-VEFS-injected (bottom) D1-MSNs in the NAc. Scale bars: 20μm. **(I)** Percentage of fluorescence intensity for H3K27me1 in GFP-positive and GFP-negative neurons. Two-way ANOVA: virus effect: F_(1,16)_=6.43; p<0.05; cell-type effect: F_(1,16)_=4.52; p<0.05; significant virus by cell-type interaction: F_(1,16)_=4.52; p<0.05. Tukey’s test: AAV-VEFS different from AAV-GFP in GFP+ cells (*p<0.05).

*In vitro* studies have demonstrated that the C-terminal domain of SUZ12, known as the VEFS-BOX domain (Figure 3D), directly interacts with EZH2 and contributes to the catalytic actions of PRC2^27, 35^, while the ΔVEFS domain determines the precise localization of the complex^38^. Strikingly, overexpression of the VEFS domain is sufficient to restore methylation of H3K27 in SUZ12 knockout cultured cells, but induces selective genome-wide enrichment of H3K27me1^32^. To determine whether the VEFS domain of SUZ12 might exert a similar effect in brain, we developed a Cre-dependent adeno-associated viral (AAV) construct expressing the VEFS domain under the control of the human synapsin (hSYN) promoter to allow for neuron-specific overexpression (AAV-VEFS, Figure 3E, Figure S3A). An AAV-hSYN-GFP construct (AAV-GFP) was used as a control. We microinfused the VEFS or GFP construct into the NAc of adult D1-Cre mice and processed tissue for molecular and neuroanatomical validations (Figure 3F). We then confirmed by qPCR that AAV-VEFS mediates increased VEFS domain expression with no change seen in levels of SUZ12 overall using primers directed outside this domain (ΔVEFS) (Figure 3G). Using immunofluorescence, we observed that AAV-VEFS increased H3K27me1 (Figure 3H-I), but not H3K27me2 or H3K27me3 (Figure S3B-E), in D1-MSNs of the NAc. Moreover, we found a significant increase in the relative abundance of H3K27me1 in bulk NAc tissue of AAV-VEFS-injected D1-Cre mice using mass spectrometry (Figure 3J). By contrast, overexpression of full-length SUZ12 did not change H3K27 methylation levels in D1-MSNs (Figure S4A-D), which indicates that the VEFS domain is sufficient to selectively induce H3K27me1 in brain *in vivo* as reported previously *in vitro*^31^.

To evaluate whether manipulating the VEFS domain confers stress-induced susceptibility, we exposed AAV-VEFS- or AAV-GFP-injected D1-Cre mice to subthreshold social defeat (SbD), which consisted of three sessions on a single day of 5 minutes of physical aggression by a novel CD-1 aggressor mouse^22^, followed by a social interaction test 24 hours later (Figure 4A). We chose SbD because this paradigm induces a rapid stress response but does not induce social avoidance in most exposed adult wild-type mice^22^. As anticipated, SbD failed to induce social avoidance in mice that received AAV-GFP, indicated by the time spent in the interaction zone in the presence of a novel CD1 mouse (social target) (Figure 4B). However, overexpression of the VEFS domain reduced the time of interaction with a CD1 social target (Figure 4B), and increased the time spent in corners of the arena in mice exposed to SbD (Figure 4C). We also found a higher percentage of susceptible mice in the AAV-VEFS group after SbD (Figure 4D).

**Figure 4.**
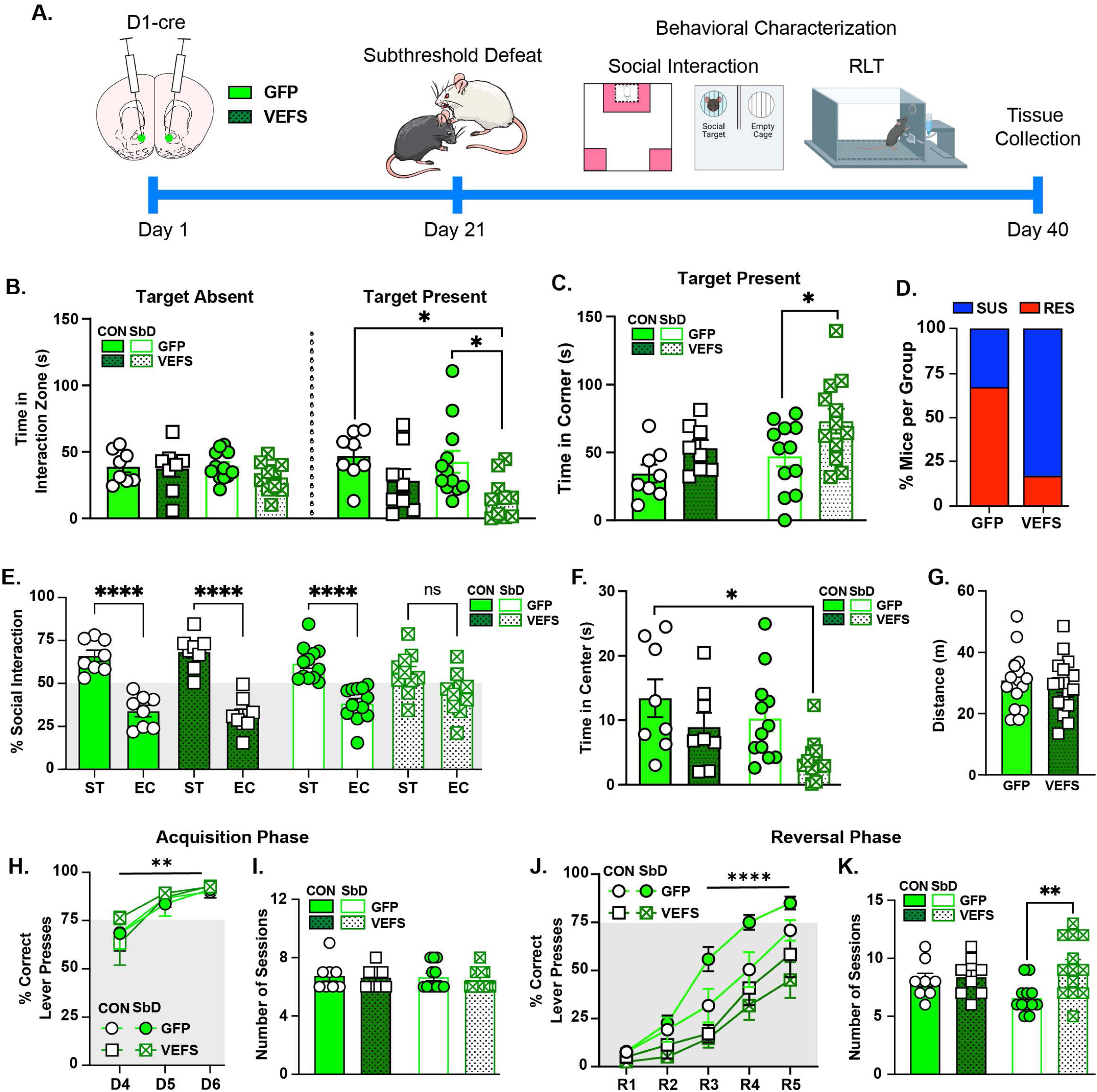
VEFS domain of SUZ12 in NAc D1-MSNs induces susceptibility to sub-threshold social defeat stress. **(A)** Timeline for GFP or VEFS viral-mediated manipulation and behavioral characterization in mice exposed to subthreshold social defeat stress (SbD) or control conditions. **(B)** Time in the social interaction zone in the absence or presence of a CD1 target. Three-way ANOVA: virus effect: F_(1,68)_=10.17; p<0.01; significant session by virus interaction: F_(1, 68)_= 4.226; p<0.05. Tukey’s test: Reduced time in the social interaction zone in the AAV-VEFS-SbD mice compared to AAV-GFP-CON and AAV-GFP-SbD groups, *p<0.05 **(C)** Time in corner. Two-way ANOVA: virus effect: F_(1,36)_=7.22; p=0.0108; group effect: F_(1,36)_=4.093; p<0.05; no significant interaction: F_(1,36)_=0.192; p=0.66. Sidak’s comparison for virus effect: Increased time in the corners in AAV-VEFS-SbD relative to AAV-GFP-SbD, *p<0.05. **(D)** Percentage of SUS and RES mice in each group. **(E)** Social interaction with a conspecific mouse in AAV-VEFS-injected mice. Three-way ANOVA: social target effect: F_(1,46)_=81.89; p<0.0001; significant social target by group interaction: F_(1,22)_=13.34; p=0.0014, significant social target by group by virus interaction: F_(1,22)_=4.57; p<0.05. Tukey’s test: Increased percentage of interaction time with social target (ST) vs empty cage (EC) in AAV-GFP-CON, AAV-VEFS-CON and AAV-GFP-SbD mice, ****p<0.0001, but not in AAV-VEFS-SbD mice: p=0.25. **(F)** Time in center of the open field. Two-way ANOVA: virus effect: F_(1,36)_=7.63; p=0.009; group effect: F_(1,36)_=4.091; p<0.05; no significant interaction: F_(1,36)_=0.209; p=0.65. Sidak’s comparison for virus effect: Decreased time in center in AAV-VEFS-SbD compared to AAV-GFP-CON, *p<0.05. **(G)** No significant differences in total distance traveled in the open field between AAV-GFP and AAV-VEFS injected mice: t_(32)_=0.59, p=0.56. **(H-I)** No significant differences between groups in the acquisition phase of the reversal learning task (RLT). **(H)** Percentage of correct lever presses: Three-way ANOVA: session effect: F_(2,111)_=18.32; p<0.0001. No significant effect of virus: F_(1,111)_=0.43; p=0.51; stress: F_(1,111)_=0.46; p=0.49, or interactions. Sidak’s comparison for session effect: D6 different from D4, *p<0.05. **(I)** Total number of sessions to reach the learning criterion: Two-way ANOVA: No significant effect of virus: F_(1,35)_=0.033; p=0.85; group: F_(1,35)_=0.51; p=0.47, or interactions. **(J-K)** VEFS overexpression led to impaired reversal learning in mice exposed to SbD. **(J**) Percentage of correct lever presses during the reversal phase: Three-way ANOVA: session effect: F_(4,180)_=64.77; p<0.0001; virus effect: F_(1,180)_=51.36; p<0.0001; group effect: F_(1,180)_=1.474; p=0.22; significant session by virus interaction: F_(4,180)_=3.084; p<0.05, significant group by virus interaction: F_(1,180)_=13.64; p<0.001. Tukey’s test: Reduced percentage of correct trials in AAV-VEFS-SbD compared to AAV-GFP-SbD, during reversal days 3 (R3), R4 and R5, ****p<0.0001. **(K)** Total number of sessions to reach the reversal learning criterion: Two-way ANOVA: virus effect: F_(1,35)_=8.16; p<0.01; group effect: F_(1,35)_=0.058; p=0.81; significant virus by group interaction: F_(1,35)_=5.88; p<0.05. Tukey’s test: Higher number of sessions in AAV-VEFS-SbD relative to AAV-GFP-SbD, **p<0.01.

We next tested whether AAV-VEFS mice similarly displayed less interaction time towards a conspecific mouse, which provides a better indication of abnormal behavior than responses to CD1 aggressors typically used in these protocols^22^. We placed experimental mice in an open field with one grid cage containing a conspecific juvenile mouse and one empty grid cage and compared the percentage of interaction time. Control mice infused with either AAV-GFP or AAV-VEFS, and SbD mice infused with AAV-GFP, displayed greater interaction towards a conspecific mouse compared to an empty cage. This social preference was lost in AAV-VEFS mice exposed to SbD (Figure 4E).

We assessed all experimental mice for exploratory behaviors, which are interpreted as being related to anxiety, using an open field test, and found that SbD mice infused with AAV-VEFS spent less time in the center area of the open field than AAV-GFP mice from the control group (Figure 4F). This effect was not associated with changes in locomotor activity since there were no differences in the total distance traveled between AAV-GFP- and AAV-VEFS-injected mice (Figure 4G).

### VEFS overexpression in NAc D1-MSNs induces stress susceptibility in more complex behavioral procedures

Mice that are susceptible to CSDS, as defined by a simple social interaction test, exhibit deficits in cognitive flexibility when assessed in an operant reversal learning task (RLT) (Figure S5A-D). Moreover, chemogenetic-induced inactivation of NAc D1-MSNs impairs this reversal learning (Figure S6). We therefore assessed whether AAV-VEFS-injected mice exposed to SbD also displayed alterations in this form of cognitive flexibility. During the first (acquisition) phase of the RLT, we trained SbD-subjected mice injected with either AAV-GFP or AAV-VEFS, and their control counterparts, to press one of two levers located on each side of the reward receptacle (active lever) to gain a 0.2% saccharin-water reward at a fixed ratio-1 (FR1) schedule. All trials were given daily until all mice reached a learning criterion of 75% correct responses during three consecutive days (Figure 4H). We did not find significant differences between groups during the acquisition phase, as all experimental mice took about the same number of sessions to achieve the learning criterion (Figure 4I). During the reversal phase, all experimental mice were trained to press the previously unrewarded lever to obtain the same 0.2% saccharin reward at a learning criterion of 75% correct responses for three consecutive days. We found that control mice injected with either AAV-GFP or AAV-VEFS exhibited similar reversal learning. By contrast, AAV-VEFS mice that were exposed to SbD displayed a reduced percentage of correct responses by reversal day 5 relative to Control-GFP mice, and by reversal days 3, 4, and 5 relative to GFP-injected SbD mice (Figure 4J). Indeed, AAV-VEFS-injected SbD mice took a greater number of sessions to complete the RLT than their AAV-GFP-injected SbD counterparts (Figure 4K), indicating that VEFS overexpression rendered mice more prone to cognitive deficits induced by a subthreshold stressor.

Finally, we assessed whether social alterations induced by VEFS overexpression can be observed in stress-naïve mice by testing a separate group of AAV-GFP- and AAV-VEFS-injected mice in an operant social reward task (Figure 5A). All experimental mice were first trained to lever press for a saccharin reward, as described above, and subsequently trained to press a lever that activated a guillotine door and gave access to sensory, but not physical, interaction with a juvenile conspecific mouse (Figure 5A). Our results show that mice from the AAV-GFP and AAV-VEFS groups displayed similar learning to lever press for a saccharin reward (Figure 5B). Interestingly, AAV-VEFS mice exhibited reduced motivation to lever press for social interaction as indicated by the lower number of breakpoints when evaluated on a progressive ratio schedule (Figure 5C).

**Figure 5.**
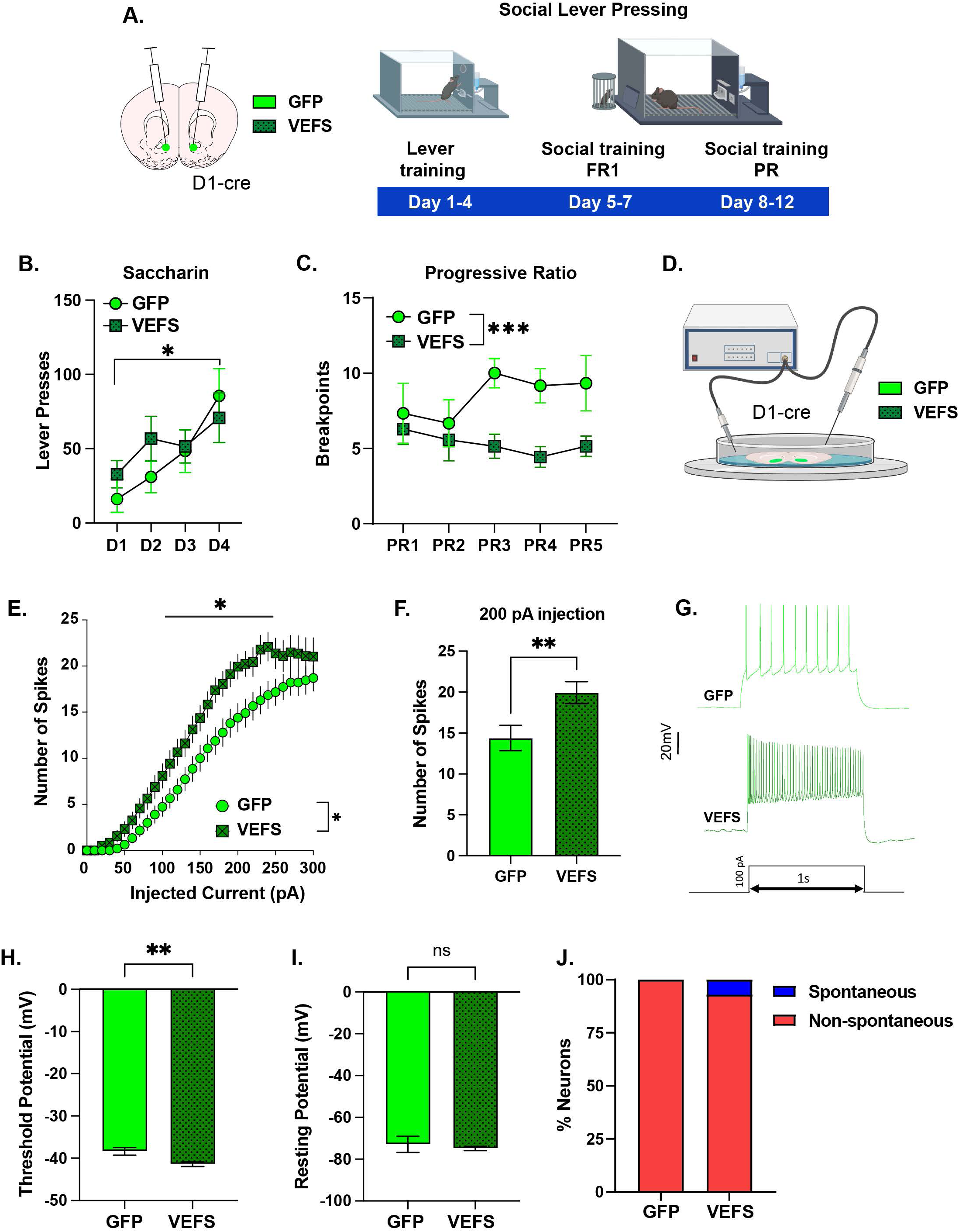
VEFS domain of SUZ12 in NAc D1-MSNs reduces social reward and enhances neuronal intrinsic excitability. **(A)** Timeline of the social reward task experiment with AAV-GFP (n=6) and AAV-VEFS (n=7). **(B)** No significant differences between groups in the acquisition of lever presses for a saccharin reward. Two-way ANOVA: session effect: F_(3,44)_=5.59, p<0.01; virus effect: F_(1,44)_=0.681, p=0.41; no significant interaction: F_(3,44)_=0.88; p=0.45. Sidak’s comparison for session effect: D5 different from D1, *p<0.05 **(C)** Progressive ratio breakpoint to lever press for a social reward. Two-way ANOVA: virus effect: F_(1,55)_=14.46; p<0.001; session effect: F_(4,55)_=0.32, p=0.85; no significant interaction: F_(4,55)_=1.09, p=0.37. Sidak’s comparison for virus effect: Lower breakpoint in AAV-VEFS-injected mice, *p<0.05. **(D)** Schematic of electrophysiology recordings in AAV-GFP- and AAV-VEFS-injected mice. **(E)** Current-evoked spikes in D1-MSNs from AAV-GFP- and AAV-VEFS-injected mice (n=41-46 neurons/6–8 mice/group). Two-way ANOVA: virus effect: F_(1,85)_=6.86; p<0.01, current effect: F_(30,2550)_=127.4; p<0.0001, significant current by virus interaction: F_(30,2550)_=1.60; p=0.02. Sidak’s comparison for virus effect: Higher number of spikes in AAV-VEFS group at 100 to 250 pA currents; *p<0.05. **(F)** Number of spikes at 200 pA current: t_(85)_= 2.713; **p<0.01. **(G)** Example voltage traces from a single D1-MSN in response to current injections. **(H)** Decreased threshold potential in AAV-VEFS-injected neurons: t_(85)_=2.95; **p<0.01. **(I)** No significant differences in the resting membrane potential of AAV-GFP- and AAV-VEFS-injected neurons: t_(85)_=0.52; p=0.59. **(J)** Percentage of D1-MSNs that exhibit spontaneous firing activity: About 8% of AAV-VEFS-injected MSNs display spontaneous firing, while no spontaneous firing was observed in GFP-injected MSNs.

Together, results from these several behavioral procedures demonstrate that a selective increase in levels of H3K27me1 in NAc D1-MSNs, as seen in susceptible mice after CSDS and induced experimentally via overexpression of the SUZ12 VEFS domain, exacerbates the emotional, social, and cognitive alterations induced by stress.

### VEFS overexpression in NAc D1-MSNs increases neuronal intrinsic excitability

Previous studies have demonstrated that stress-susceptible mice display enhanced intrinsic excitability of D1-MSNs of the NAc^13, 15, 39^. Therefore, we hypothesized that VEFS overexpression would mimic stress-induced electrophysiological alterations of this cell type. We injected stress-naïve D1-Cre mice with either AAV-GFP or AAV-VEFS into the NAc, and conducted whole-cell patch clamp recordings to examine the intrinsic excitability of fluorescently-tagged neurons (Figure 5D). As hypothesized, we observed a dramatic increase in current-induced neuronal firing in AAV-VEFS-injected mice at 100 to 250 pA relative to AAV-GFP-injected mice (Figure 5E-G), an effect that associated with a decreased threshold to fire an action potential (Figure 5H), but with no change in resting membrane potential (Figure 5I). We also observed a small percentage (∼8%) of AAV-VEFS-injected MSNs that fired spontaneously, while none of the AAV-GFP-injected MSNs exhibited spontaneous firing (Figure 5J)—consistent with the known lack of spontaneous firing of normal D1-MSNs in brain slices. By contrast, no overt behavioral or electrophysiological effects were observed following overexpression of the full-length SUZ12 (Figure S4E-I)

### ELS induces a persistent increase in H3K27me1 levels in the NAc

To evaluate whether changes in H3K27 methylation are observed across stress models and could serve as an epigenetic scar, we focused on ELS, a paradigm known to heighten stress vulnerability in adulthood^2, 6–8^. We used a recently generated mass spectrometry dataset of histone profiling in NAc tissue from mice exposed to ELS or their standard-raised (Std) counterparts^6^. In this study, ELS was induced by exposing pups to the combination of maternal separation and reduced nesting material during postnatal days (PND) 10 to PND17, and tissue was collected at PND21 and in adulthood (Figure 6A). Consistent with our results in CSDS-susceptible mice, we observed a significant increase in H3K27me1 abundance in the NAc of mice exposed to ELS. This effect was observed at PND21, immediately after ELS exposure, and lasted into adulthood, when the abundance of H3K27me1 is normally very low in mice raised under standard conditions (Figure 6B). By contrast, we observed a significant decrease in the abundance of H3K27me2 at PND21 only (Figure 6C). These findings confirm our hypothesis that different forms of stress can induce accumulation of H3K27me1 in NAc and that these effects can persist long after stressors have ceased.

**Figure 6.**
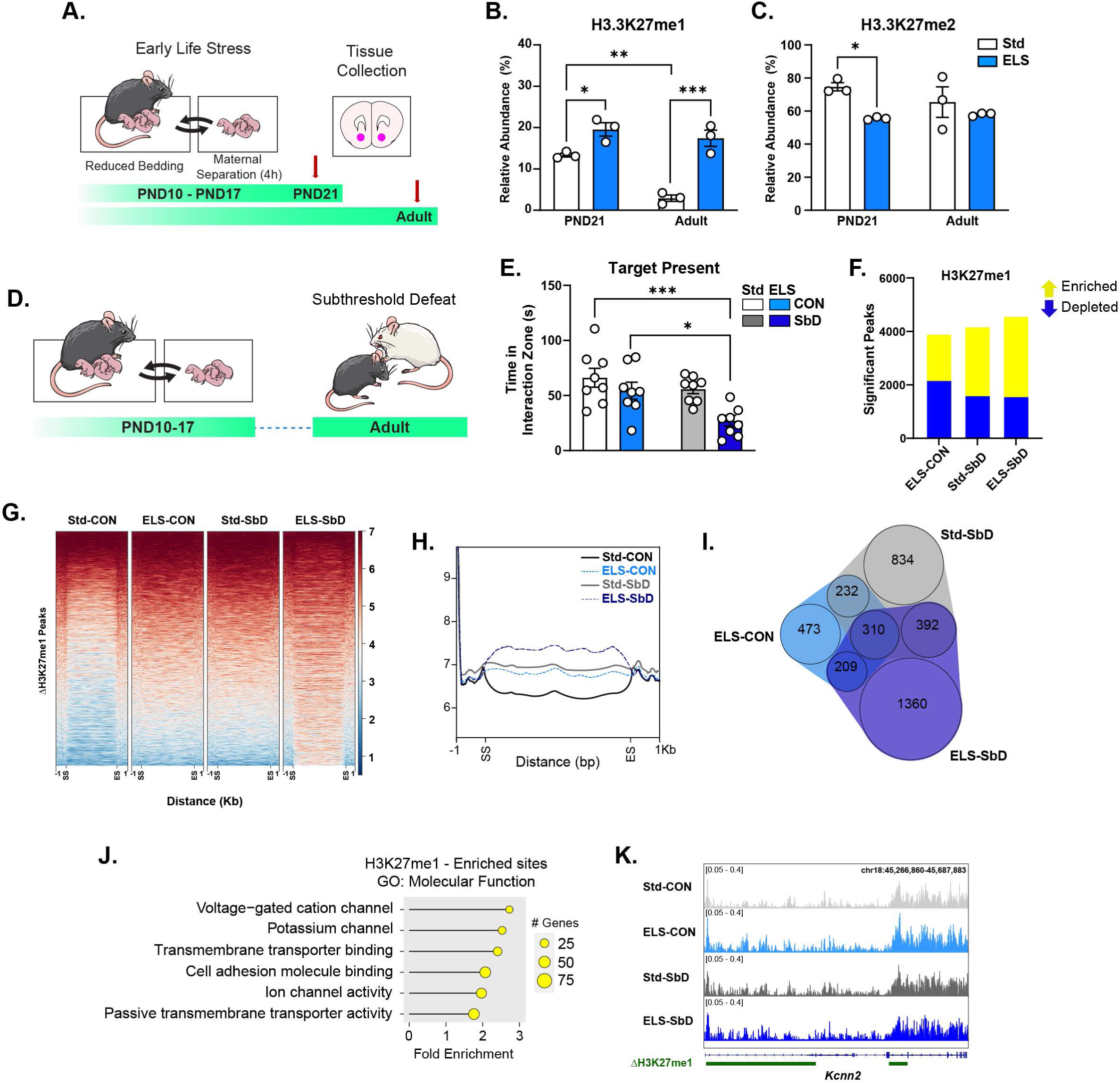
Persistent increase in H3K27me1 in NAc after ELS. **(A)** Timeline of the ELS experiment. Pups from the ELS group are subjected to maternal separation and reduced bedding in their home-cage during PND10-17. NAc tissue from mice exposed to ELS and standard-raised condition (Std) was collected at PND21 and adulthood (∼8 weeks). **(B)** ELS increases the relative abundance of H3K27me1 across the lifespan. Two-way ANOVA: ELS effect: F_(1,8)_=59.11; p<0.0001; age effect: F_(1,8)_=21.7; p<0.01; significant ELS by age interaction: F_(1,8)_=9.349; p<0.05. Tukey’s test: Increased H3K27me1 in the ELS group at PND21 (*p<0.05) and Adult (***p<0.001), compared to Std group. Decreased H3K27me1 in the Std group in Adult, compared to Std group at PND21, **p<0.01. **(C)** Transient reduction of H3K27me2 after ELS. Two-way ANOVA: ELS effect: F_(1,8)_=7.675; p<0.05. No significant age effect: F_(1,8)_=0.403, p=0.54; or interactions: F_(1,8)_=1.67, p=0.23. Sidak’s comparison for ELS effect: ELS group different from Std group at PND21, *p<0.05. **(D)** Timeline of the “double-hit” paradigm: pups were exposed to ELS and to a subsequent exposure to subthreshold social defeat (SbD) in adulthood (n=8/group). **(E)** Reduced social interaction in mice exposed to both ELS and SbD. Two-way ANOVA: ELS effect: F_(1,28)_=10.19, p<0.01; SbD effect: F_(1,28)_=8.78, p<0.01; ELS by SbD interaction: F_(1,28)_=1.94, p=0.17. Sidak’s comparison for ELS effect: ELS-SbD group different from Std-CON (***p<0.001), and ELS-CON (*p<0.05). **(F)** Number of differential enriched (yellow) of depleted (blue) peaks for H3K27me1 in the ELS-CON vs Std-CON, Std-SbD vs Std-CON, and ELS-SbD vs Std-CON comparisons. **(G)** Coverage heatmap for H3K27me1 enrichment within a region spanning ± 1 kb around the start site (SS) and end site (ES) of the differential peaks in the ELS-SbD vs Std-CON comparison. The gradient blue-to-red color indicates low-to-high counts. **(H)** Average H3K27me1 density within differential (ELS-SbD vs Std-CON) peaks and their 1 kb flanking regions. **(I)** Venn diagram of differential H3K27me1 peaks in Std-CON, ELS-CON and ELS-SbD vs Std-CON comparisons. **(J)** Top 6 GO molecular function terms associated with H3K27me1 enrichment in the ELS-SbD vs Std-CON comparison. Terms were ranked according to their log-10(FRD), followed by fold enrichment. **(K)** Representative IGV browser tracks of H3K27me1 enrichment across the potassium channel gene, *Kcnn2*. ΔH3K27me1 sites span across introns 1 and 3.

We next examined whether ELS alters H3K27me1 deposition across the genome, and how this deposition is affected by a second hit of adult stress (Figure 6D). Pups were exposed to either ELS or Std conditions, as described above, and allowed to reach adulthood, when a subset of these mice was subjected to a second hit of adult SbD stress. NAc tissue was collected 24 hours after the SIT and then processed for CUT&RUN-seq for H3K27me1. As expected, adult mice exposed to ELS alone or to SbD alone did not exhibit social avoidance, however, the combination of ELS and SbD reduced social interaction time in comparison to other conditions (Figure 6E), which indicates that ELS heightens vulnerability not only to adult exposure to chronic stress^6–8^ but also to subthreshold stressors. Moreover, the “double-hit” stress impaired cognitive flexibility in the RLT in an independent cohort of mice (Figure S5E-G).

We analyzed the differentially-enriched sites for H3K27me1 in the NAc of ELS-control, Std-SbD and ELS-SbD conditions compared to the Std-control group. This approach revealed the largest number of differential events in the ELS-SbD vs Std-control comparison, and about the same number of peaks in mice exposed to ELS-control vs Std-Control, and Std-SbD vs Std-Control, comparisons (Figure 6F). Therefore, exposure to ELS or SbD alone was insufficient to induce dramatic enrichment of H3K27me1, but the combination of the two stressors was effective at doing so. This finding was further confirmed by heatmaps, which showed the strongest H3K27me1 peak signals in the ELS-SbD vs Std-control comparison (Figure 6G-H). As shown by Venn diagrams, the ELS-SbD vs Std-control comparison exhibited the greatest number of differentially-enriched sites, with some overlap of H3K27me1-enriched sites among the various conditions (Figure 6I).

Finally, we applied GO analysis to the genes that displayed altered H3K27me1 deposition in the ELS-SbD vs Std-control comparison and identified molecular terms associated with neuronal activity (Figure 6J, Table S3), such as “*Voltage-gated cation channel activity*”, or “*Potassium channel activity*”, which indicates once again that genes involved in maintaining excitability of MSNs are a preferential target for H3K27me1 regulation in the NAc (Figure 6K).

## DISCUSSION

Various methylated states of H3K27 correspond to >80% of total H3, while the unmodified or acetylated forms of H3K27 are restricted to only about 15% and 2%, respectively^31, 40^. H3K27me3 has been widely explored and consistently associated with gene repression due to its preferential accumulation across heterochromatin sites and at promoter regions of silenced genes^21, 41^. By contrast, the roles of H3K27me1 and H3K27me2 have been poorly studied in the brain despite their higher occurrence compared to H3K27me3^31^, and little is known about the mechanism by which these marks influence transcriptional states, either across development or in response to environmental challenges. Here, we identified H3K27me1 as the most prominent histone modification induced in the NAc by CSDS, and found that it was likewise induced in a persistent manner by ELS. CSDS and ELS are two extensively-characterized chronic stress models in mice that induce long-lasting behavioral abnormalities^7, 23^. Following an unbiased proteomics strategy, we found increased H3K27me1 abundance in: *(i)* adult mice that are susceptible to CSDS (Figure 1), and *(ii)* juvenile and adult mice previously exposed to ELS (Figure 6). In both cases, the increase in H3K27me1 was associated with decreased abundance of H3K27me2, a striking finding given that the two marks exist at mostly distinct genomic regions. On a genome-wide scale, H3K27me1 is enriched at intragenic and promoter regions, particularly at genes associated with neuronal excitability. Moreover, using immunofluorescence and viral-mediated manipulations, we demonstrated that overexpressing the VEFS domain, the catalytic moiety of SUZ12, selectively increased H3K27me1 levels in NAc D1-MSNs, which enabled us to directly implicate this lasting histone modification in the behavioral abnormalities induced by stress and in altered intrinsic neuronal excitability. Our highly novel findings support the hypothesis that H3K27me1 in NAc D1-MSNs represents a “chromatin scar” that mediates stress susceptibility across the lifespan.

Evidence from *in vitro* studies shows that the effect of H3K27me1 on gene expression is complex, and depends on its precise location within genomic regions or the cell type where it is enriched^27, 31, 42, 43^. For example, H3K27me1 is deposited over intragenic regions of active genes in neuroblastoma cells^43^, lymphocytes^44^ and mouse embryonic stem cells^27, 31^, whereas depletion of this mark is observed in the promoters of actively transcribed genes in HeLa cells^42^. Using bulk NAc tissue, we generated the first dataset of genome-wide enrichment of H3K27me1 and H3K27me2 in adult brain, and confirmed previous *in vitro* observations reporting the preferential deposition of H3K27me1 within gene bodies and promoters, and of H3K27me2 across intergenic regions. Our CUT&RUN-seq experiments further revealed that mice susceptible to CSDS displayed a larger number of differential H3K27me1-enriched sites relative to resilient mice (Figure 2B). A similar observation was found in our ELS dataset, where the largest number of enhanced H3K27me1 peaks was seen in mice exposed to ELS and then to a second stress in adulthood (Figure 6H). The top molecular function of genes that exhibited higher H3K27me1 deposition in response to either stress related to neuronal excitability, such as GABA receptor subunits or potassium channels, both of which have consistently been linked to depression and anxiety^13, 45–48^. We also found that CSDS led to depletion of H3K27me1 within intragenic and promoter sites of genes involved with histone methyltransferase activity and overall RNA regulation (Figure 2I). These findings suggest that stress alters the pattern of H3K27me1 accumulation within particular genomic regions and thereby influences different molecular functions. This may indicate that specific H3K27me1-enriched sites are more accessible to the activity of PRC2 and, in turn, may bias its regulatory actions towards those sites at the expense of other regions^32, 37^.

Work in cultured cells has shown that the pattern of H3K27 methylation can be shaped by the enrichment of SUZ12 at precise target regions, a process mediated by the C-terminal VEFS and N-terminal (ΔVEFS) domains^27, 29, 32, 38, 49^. Thus, while the VEFS domain is sufficient to recruit PRC2 and catalyze H3K27 methylation, the ΔVEFS domain determines the precise localization of SUZ12 by recognizing specific genomic regions. Furthermore, overexpression of the VEFS domain, but not ΔVEFS, restores all three H3K27 methylation states in SUZ12 knockout cells, but induces aberrant accumulation of H3K27me1 only across active gene bodies^32, 36, 50^. We took advantage of this *in vitro* observation and, using viral-mediated manipulations, revealed that VEFS overexpression elevated levels of H3K27me1, but not H3K27me2 or H3K27me3, in D1-MSNs of the NAc. This manipulation induced behavioral and cellular signatures of stress susceptibility, such as increased social avoidance, impaired cognitive flexibility, and enhanced intrinsic excitability of D1-MSNs^13, 15, 16, 39, 51^. Moreover, this effect was specific to the VEFS domain, since we did not observe stress-related phenotypes by overexpressing full-length SUZ12 (Figure S4). Our observation that CSDS induces SUZ12 in NAc, but that SUZ12 overexpression does not per se yield the stress-induced pattern of H3K27me1 regulation or downstream behavioral abnormalities, suggests that there are other factors operating in D1-MSNs of stressed mice to induce the molecular, cellular and behavioral phenotypes observed. An improved understanding of PRC2 functioning in brain, and identification of the rich regulatory mechanisms that control its activity and genomic targeting, will be required to identify such additional stress-related factors.

Robust evidence demonstrates that D1-MSNs and D2 MSNs display distinct molecular and electrophysiological signatures and contribute differentially to stress-related phenotypes^15, 16, 39^. Our previous work has shown that stress occurring in early life shapes the transcriptional profiles of the brain’s reward circuitry^6–8^. This transcriptional regulation is mediated, in part, by enrichment of H3K79me2 in D2 MSNs of the NAc, and the coordinated action of its writer, DOT1L, and eraser, KDM2B, enzymes^6^. Using the same mass spectrometry dataset as in^6^, we identified H3K27me1 as a persistent histone modifications induced by ELS (Figure 6), which, along with our proteomics, RNAscope and viral-manipulation experiments, indicate that these changes predominate in D1-MSNs. Despite the inability to obtain cell-specific histone proteomics data due to the amount of tissue required, we hypothesize that stress induces these two distinct histone modifications in these two distinct MSN cell types and thereby causes cell-type-specific transcriptional and physiological regulation. Furthermore, given that we did not observe changes in H3K79me2 in adult mice exposed to CSDS (Figure S1), we also argue that stress-induced changes in histone modifications are age-dependent and highly sensitive to developmental windows of brain plasticity. In this context, certain histone marks may be more likely to be enriched or depleted with high spatial and temporal specificity as compared to others. Much remains unknown about whether stress-induced changes of H3K27me1 in D1-MSNs may lead to specific crosstalk between histone modifications^52^, thus, facilitating the generation or depletion of other marks, and, as a result, altering the epigenetic landscape in other cell types of the NAc. This is a provocative question given previous reports in cultured cells showing that, in the absence of H3K27me2, there is aberrant deposition of H3K27me1 at intragenic regions of poorly expressed genes and a global, but diffuse, increase of H3K27ac accumulation along the genome^31, 36^. The fact that stress also reduced the accumulation of H3K27me1 within genes involved in histone methyltransferase activity may support this idea.

In summary, by combining unbiased proteomics profiling, genome-wide approaches, and mouse stress models, our work establishes a novel function of H3K27me1 in the NAc in serving as a chromatin scar that mediates lifelong stress susceptibility. We report that accumulation of H3K27me1 in NAc D1-MSNs leads to persistent emotional, social and cognitive abnormalities, as well as enhanced excitability of these neurons, and we map genome-wide targets for this mark that now require greater investigation as mediators of these phenotypes. An important technical contribution of our work is to introduce the catalytic VEFS domain of SUZ12 as driving the aberrant deposition of H3K27me1 in response to several forms of stress. Together, our findings thus open the door for several new therapeutic targets for human stress disorders.

## ACKNOWLEDGEMENTS

This work was supposed by grants from the National Institute of Mental Health (R01MH051399 and R01MH129306 to EJN), the Hope for Depression Research Foundation (to EJN), and the Robin Chemers Neustein Postdoc Award and the FBI Postdoc Innovation Award (to ATB). The Sidoli lab gratefully acknowledges for funding the Einstein-Mount Sinai Diabetes center, Merck, Relay Therapeutics, Deerfield (Xseed award) and the NIH Office of the Director (S10OD030286). We thank Dr. Kristian Helin (The Institute of Cancer Research, London) for kindly providing the SUZ12 and VEFS plasmids, as well as Dr. Ramesh Chandra from the Virus Vector Core of the University of Maryland School of Medicine for packing the viral constructs. This study used the services of Mount Sinai’s electrophysiology core, under the direction of Dr. Robert D. Blitzer. An earlier version of this manuscript was posted on bioRxiv.

## AUTHOR CONTRIBUTIONS

A.T.B. and E.J.N. designed the studies. A.T.B. performed behavioral experiments, and processed samples for histone isolation, library preparations, and neuroanatomical analysis. M.E. and A.R. performed the sequencing-based bioinformatics with input from L.S. O.I. and A.T.B. amplified the plasmids. A.T.B., O.I., A.M.-T., R.D.d.C., C.J.B., E.M.P., Y.v.d.Z., D.W., F.J., M.-R. performed stereotaxic surgeries and contributed to tissue collection. H.K. provided the mass spectrometry dataset for the ELS study. V.P. conducted electrophysiology experiments. S.S. conducted mass spectrometry experiments. A.T.B. and E.J.N. wrote the manuscript. E.J.N. supervised the project. All authors discussed the results and commented on and edited the paper.

## DATA AND CODE AVAILABILITY

CUT&RUN-seq data is deposited at GEO (Pending number) and will be publicly available as of the date of publication. This study did not generate original code.

