## Supplemental Information for "Monomethylation of Lysine 27 at Histone 3 Confers Lifelong Susceptibility to Stress"

### Figure S1

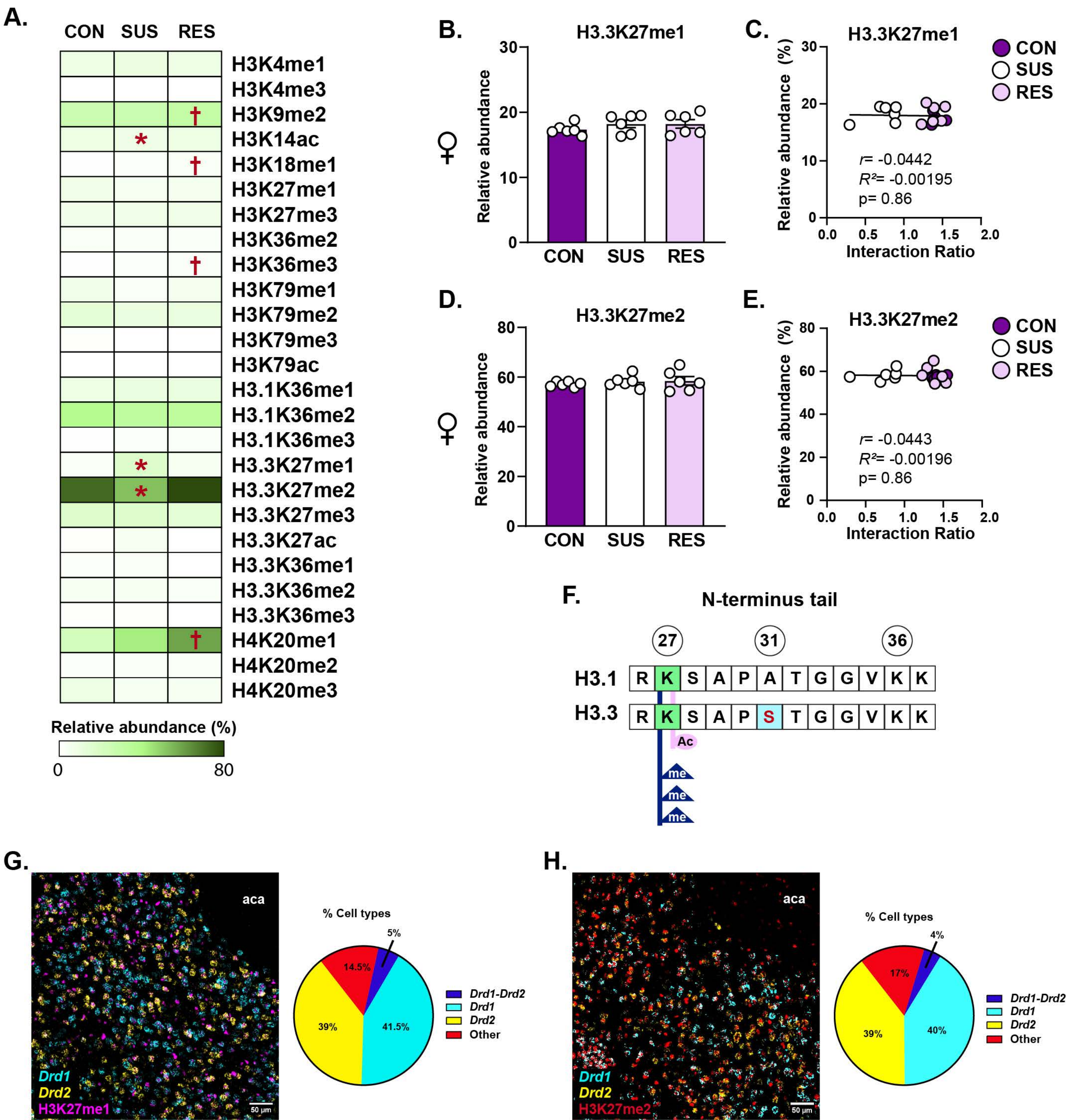
A.
CON SUS RES
H3K4me1
H3K4me3
H3K9me2
H3K14ac
H3K18me1
H3K27me1
H3K27me3
H3K36me2
H3K36me3
H3K79me1
H3K79me2
H3K79me3
H3K79ac
H3.1K36me1
H3.1K36me2
H3.1K36me3
H3.3K27me1
H3.3K27me2
H3.3K27me3
H3.3K27ac
H3.3K36me1
H3.3K36me2
H3.3K36me3
H4K20me1
H4K20me2
H4K20me3
Relative abundance (%)
0
80
B.
H3.3K27me1
Relative abundance
CON SUS RES
C.
H3.3K27me1
Relative abundance (%)
Interaction Ratio
CON
SUS
RES
r= -0.0442
R²= -0.00195
p= 0.86
D.
H3.3K27me2
Relative abundance
CON SUS RES
E.
H3.3K27me2
Relative abundance (%)
Interaction Ratio
CON
SUS
RES
r= -0.0443
R²= -0.00196
p= 0.86
F.
N-terminus tail
27 31 36
H3.1 R K S A P A T G G V K K
H3.3 R K S A P S T G G V K K
Ac
me
me
me
G.
aca
Drd1
Drd2
H3K27me1
50 µm
% Cell types
5%
14.5%
39%
41.5%
Drd1-Drd2
Drd1
Drd2
Other
H.
aca
Drd1
Drd2
H3K27me2
50 µm
% Cell types
4%
17%
39%
40%
Drd1-Drd2
Drd1
Drd2
Other

### Figure S2

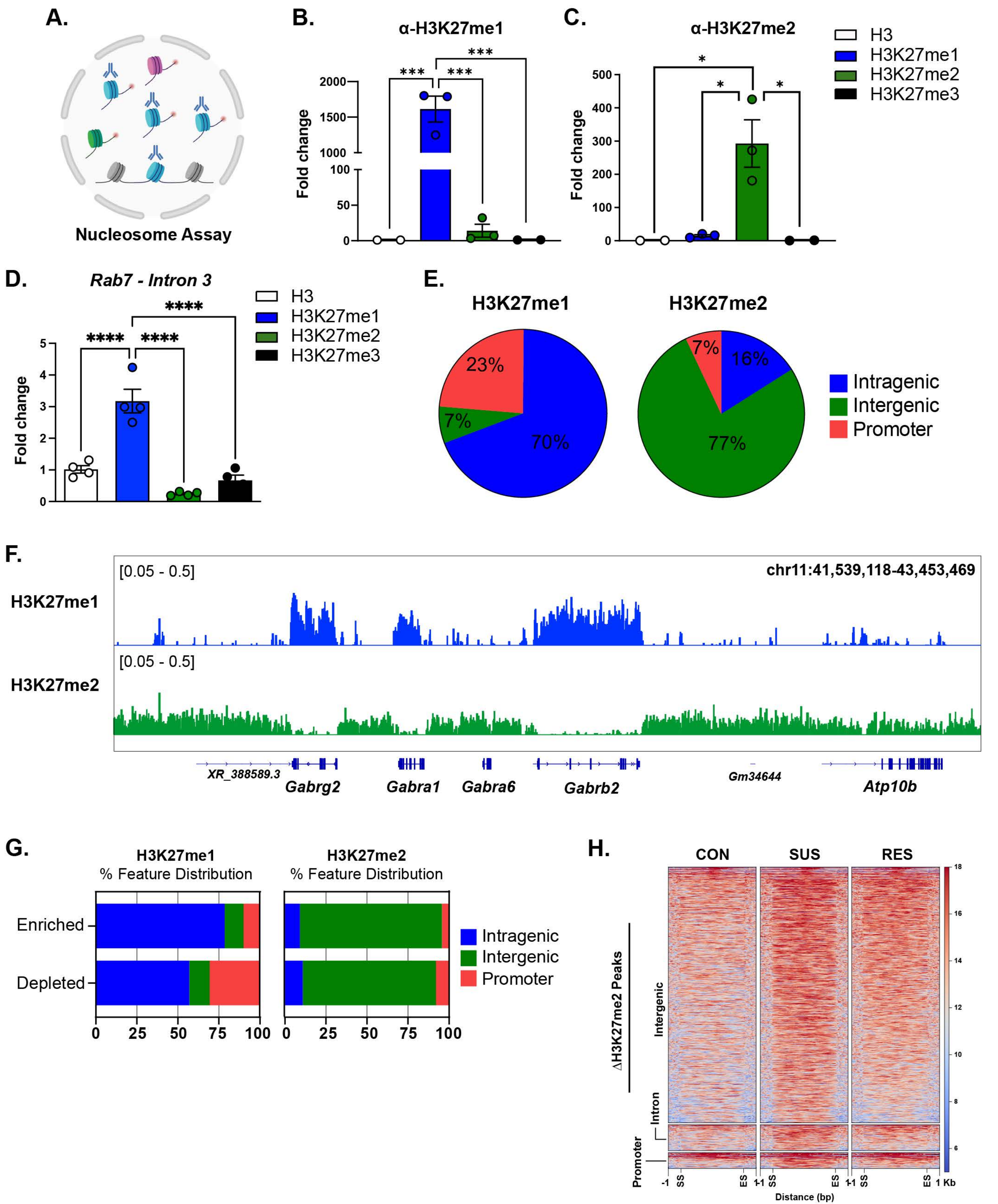

A.
Nucleosome Assay
B.
α-H3K27me1
Fold change
C.
α-H3K27me2
H3
H3K27me1
H3K27me2
H3K27me3
D.
*Rab7 - Intron 3*
E.
H3K27me1
H3K27me2
23%
7%
70%
7%
16%
77%
Intragenic
Intergenic
Promoter
F.
[0.05 - 0.5]
chr11:41,539,118-43,453,469
XR_388589.3
Gabrg2
Gabra1
Gabra6
Gabrb2
Gm34644
Atp10b
G.
% Feature Distribution
Enriched
Depleted
H.
CON
SUS
RES
ΔH3K27me2 Peaks
Intergenic
Intron
Promoter
Distance (bp)

### Figure S3

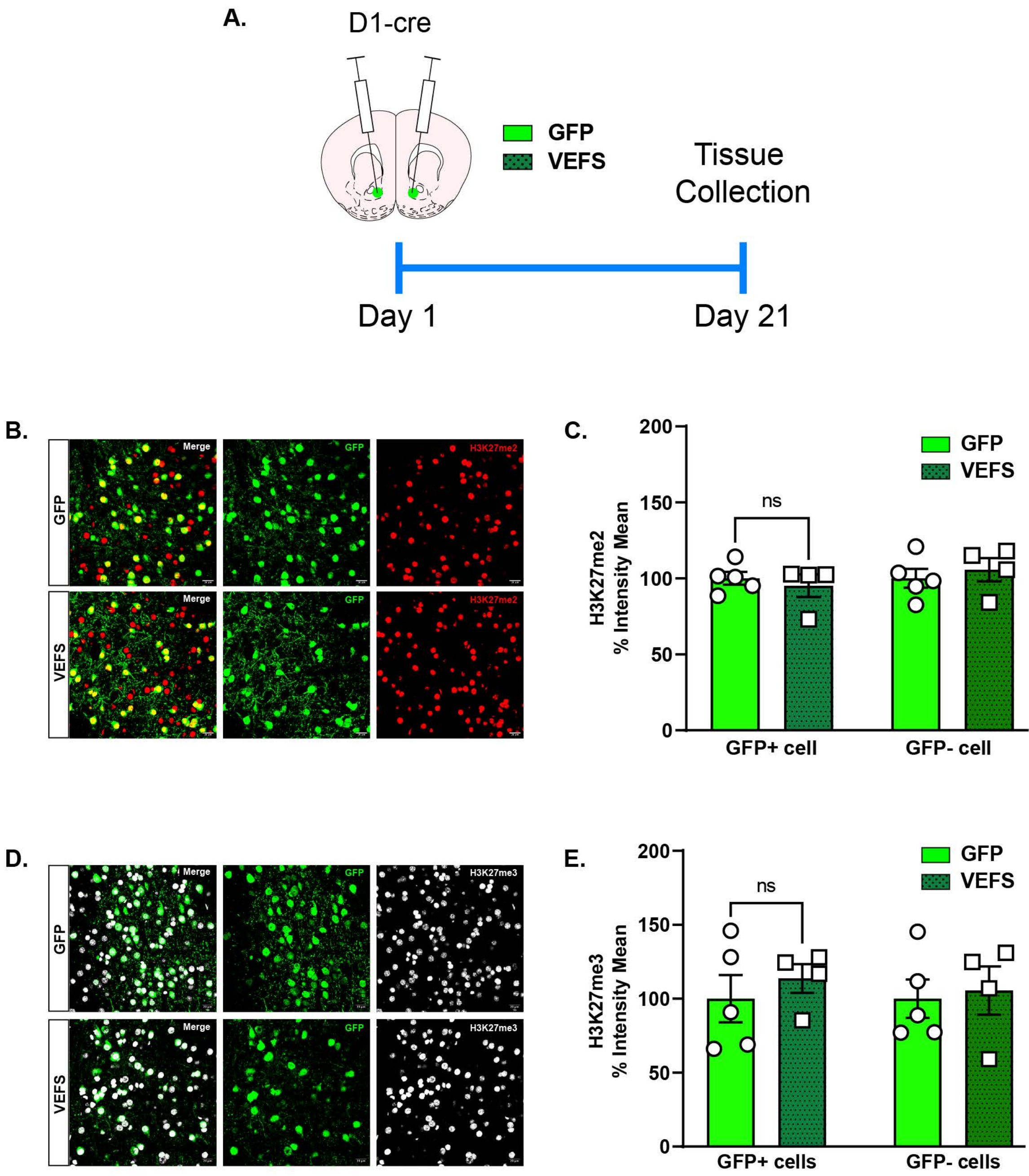
A.
D1-cre
GFP
VEFS
Tissue Collection
Day 1
Day 21
B.
Merge
GFP
H3K27me2
GFP
VEFS
C.
200
150
100
50
0
H3K27me2 % Intensity Mean
ns
GFP
VEFS
GFP+ cell
GFP- cell
D.
Merge
GFP
H3K27me3
GFP
VEFS
E.
200
150
100
50
0
H3K27me3 % Intensity Mean
ns
GFP
VEFS
GFP+ cells
GFP- cells

### Figure S4

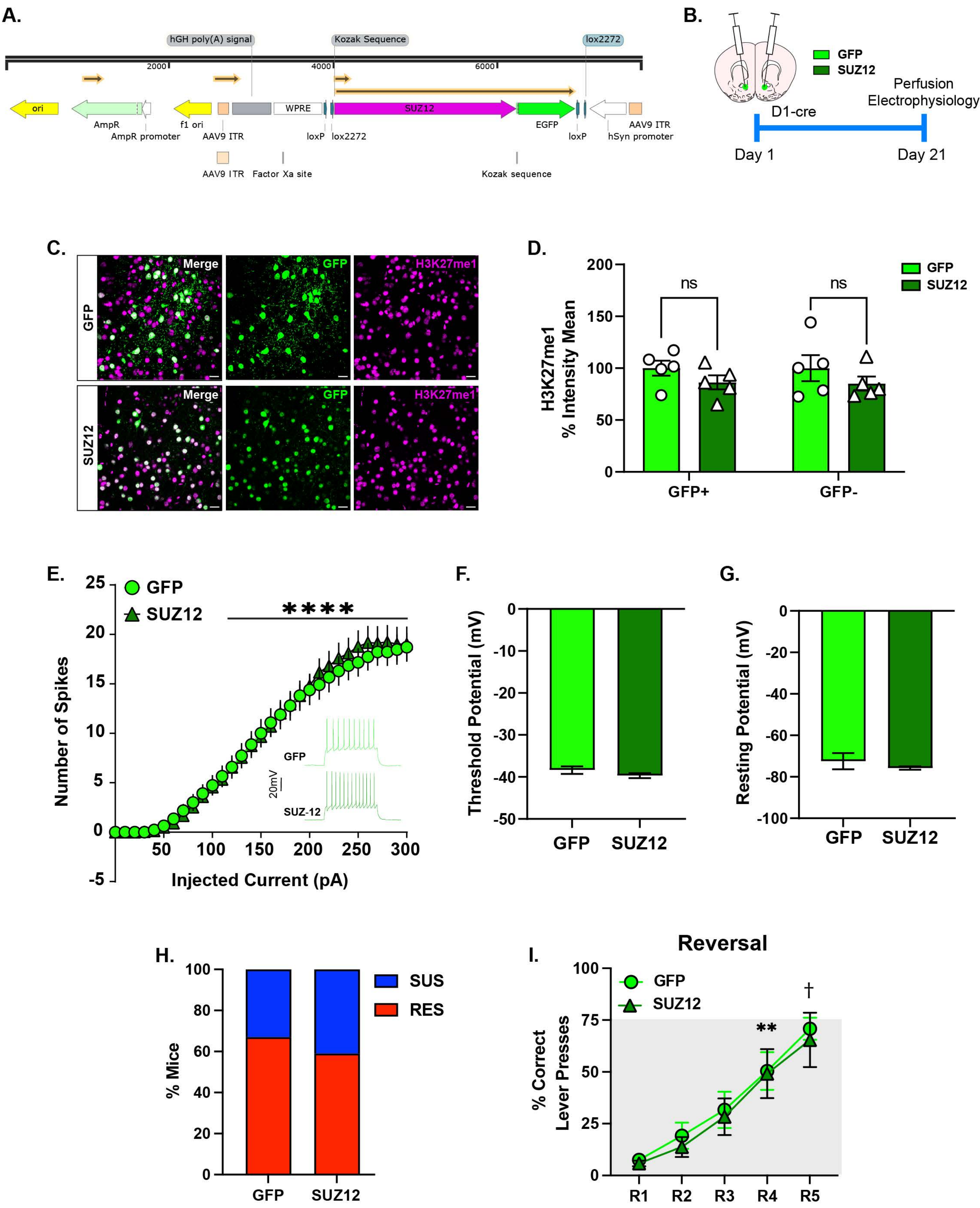

A.
hGH poly(A) signal
Kozak Sequence
lox2272
2000
4000
6000
ori
AmpR
AmpR promoter
f1 ori
AAV9 ITR
WPRE
loxP
lox2272
SUZ12
EGFP
loxP
hSyn promoter
AAV9 ITR
AAV9 ITR
Factor Xa site
Kozak sequence
B.
GFP
SUZ12
D1-cre
Perfusion
Electrophysiology
Day 1
Day 21
C.
Merge
GFP
H3K27me1
GFP
SUZ12
D.
H3K27me1
% Intensity Mean
GFP
SUZ12
ns
ns
GFP+
GFP-
E.
GFP
SUZ12
****
Number of Spikes
Injected Current (pA)
GFP
SUZ-12
20mV
F.
Threshold Potential (mV)
GFP
SUZ12
G.
Resting Potential (mV)
GFP
SUZ12
H.
% Mice
SUS
RES
GFP
SUZ12
I.
Reversal
% Correct
Lever Presses
GFP
SUZ12
**
†
R1
R2
R3
R4
R5

### Figure S5

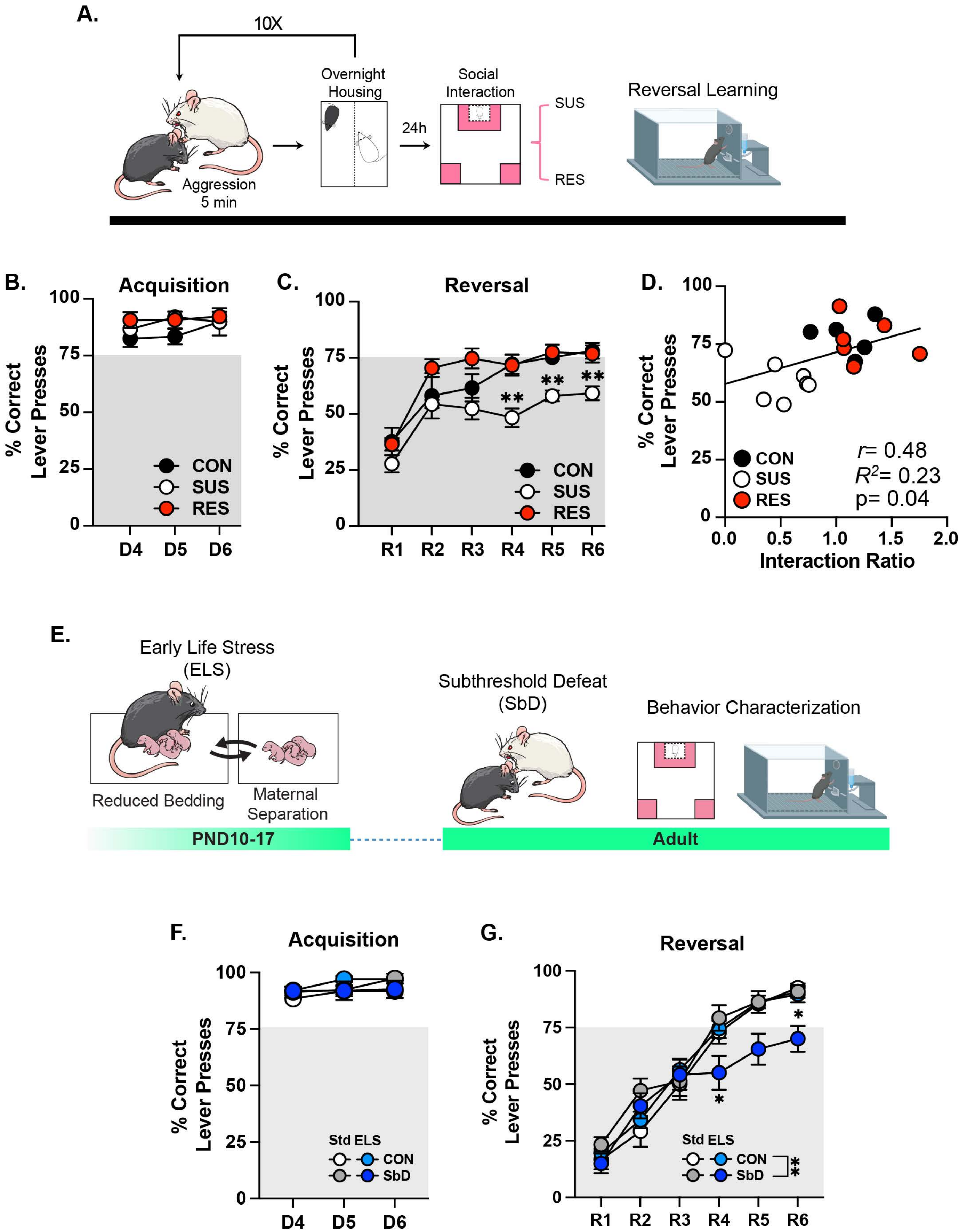

A.
10X
Overnight Housing
24h
Social Interaction
SUS
RES
Reversal Learning
Aggression 5 min
B.
Acquisition
% Correct Lever Presses
CON
SUS
RES
D4 D5 D6
C.
Reversal
R1 R2 R3 R4 R5 R6
D.
r= 0.48
R2= 0.23
p= 0.04
Interaction Ratio
E.
Early Life Stress (ELS)
Reduced Bedding
Maternal Separation
Subthreshold Defeat (SbD)
Behavior Characterization
PND10-17
Adult
F.
Acquisition
Std ELS
CON
SbD
G.
Reversal

### Figure S6

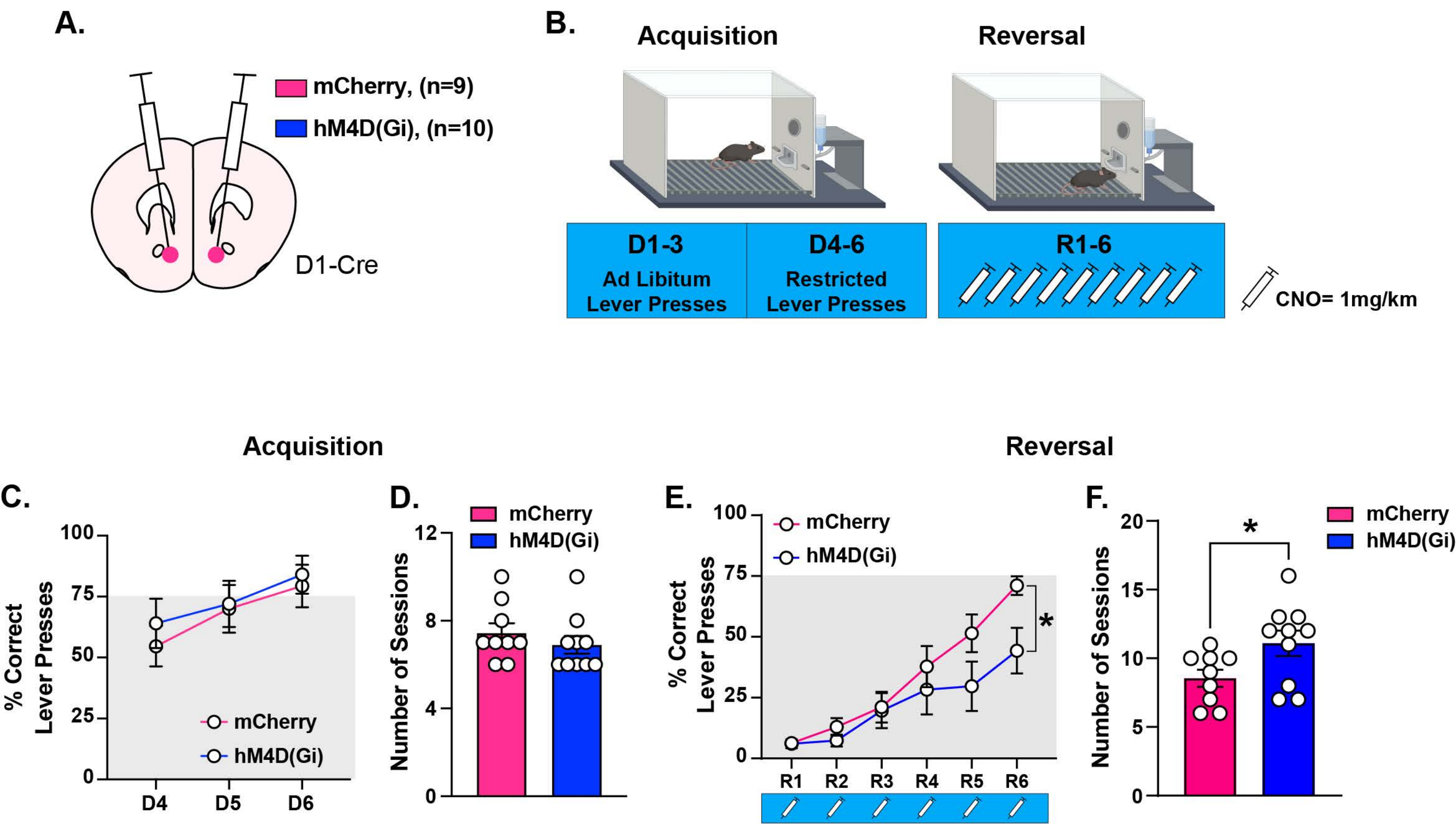
A.
mCherry, (n=9)
hM4D(Gi), (n=10)
D1-Cre
B.
Acquisition
Reversal
D1-3
Ad Libitum
Lever Presses
D4-6
Restricted
Lever Presses
R1-6
CNO= 1mg/km
Acquisition
Reversal
C.
% Correct
Lever Presses
mCherry
hM4D(Gi)
D4
D5
D6
D.
Number of Sessions
mCherry
hM4D(Gi)
E.
% Correct
Lever Presses
mCherry
hM4D(Gi)
R1
R2
R3
R4
R5
R6
*
F.
Number of Sessions
mCherry
hM4D(Gi)
*

#### SUPPLEMENTAL FIGURE LEGENDS

**Figure S1. Relative abundance of histone modifications in NAc after CSDS. (A)** Heatmap displays relative abundance obtained for the modifications shown on H3 and H4 K residues in control (CON, n=6), susceptible (SUS, n=5) and resilient (RES, n=5) mice. Mono/di/trimethylation – me1/2/3; acetylation – ac. One-way ANOVA of each mark was calculated. Tukey's test: SUS different from CON and RES, *$p<0.0001$; RES different from CON, †$p<0.05$. **(B)** Relative abundance of H3.3K27me1 in females after CSDS: One-way ANOVA: $F_{(2,15)}=0.85$; $p=0.44$. **(C)** Lack of correlation between the relative abundance of H3K27me1 in NAc and the social interaction ratio. **(D)** Relative abundance of H3K27me2 in females after CSDS: One-way ANOVA: $F_{(2,15)}=0.37$; $p=0.69$. **(E)** Lack of correlation between the relative abundance of H3.3K27me2 in NAc and the social interaction ratio. **(F)** Depiction of the N-terminus corresponding to H3.1 and H3.3 variants, and the K27, Alanine (A) 31, and K36 residues. Histone H3.3 contains a serine (S) for alanine (A) substitution at position 31 (S31, in blue). Antibodies for acetylation or methylation of K27 (in green) recognize these histone modifications in H3.1 and H3.3. **(G)** Left panel: Representative coronal section showing H3K27me1 expression in *Drd1* (D1) and *Drd2* (D2) MSNs of the NAc in adult stress-naïve mice. Right panel: Percentage of H3K27me1 expression by cell-type. **(H)** Left panel: H3K27me2 expression in D1- and D2-MSNs. Right panel: Percentage of H3K27me2 expression by cell-type. Scale bar: 20µm.

**Figure S2. Validation of CUT&RUN for H3K27me1 and H3K27me2. (A)** Nucleosome assay. Chromatin was added with purified recombinant nucleosomes and incubated with antibodies for either H3K27me1 or H3K27me2. Each recombinant nucleosome contained one of the methylation forms of K27 and could be distinguished by a unique DNA sequence (“barcode”). Following DNA purification, each barcode was amplified with specific primers using qPCR. **(B)** Specific amplification with H3K27me1 primers in chromatin incubated with H3K27me1 antibody. One-way ANOVA: $F_{(3,6)}=54.09$; $p<0.0001$. Tukey's test: ***Different from H3, H3K27me2 and H3K27me3, $p<0.001$. **(C)** Specific amplification with H3K27me2 primers in chromatin incubated with H3K27me2 antibody. One-way ANOVA: $F_{(3,6)}=11.21$; $p=0.01$. Tukey's test: *Different from H3, H3K27me1 and H3K27me3, $p<0.05$. **(D)** CUT&RUN followed by qPCR of NAc tissue incubated with antibodies for total H3 and for the three H3K27 methylation forms. Selective amplification of the H3K27me1-enriched gene, *Rab7*. One-way ANOVA: $F_{(3,12)}=37.70$; $p<0.0001$. Tukey's test: *Different from H3, H3K27me2 and H3K27me3, ****$p<0.0001$. **(E)** Feature distribution of H3K27me1 and H3K27me2 enrichment under basal condition. **(F)** Representative Integrative Genomics Viewer (IGV) browser tracks showing H3K27me1 (in blue) and H3K27me2 (in green) enrichment over different $GABA_A$ receptor associated genes across chromosome 11. **(G)** Feature distribution of enrichment and depletion of H3K27me1 and H3K27me2 in the SUS vs CON comparison. H3K27me1 deposition occurs mainly over intragenic regions, whereas H3K27me2 is restricted to intergenic regions**.** (H) Coverage heatmap for H3K27me2 enrichment within a region spanning ± 1 kb around the start site (SS) and end site (ES) of the differential peaks in the SUS vs CON comparison. The gradient blue-to-red color indicates low-to-high counts within intragenic, promoter, exon and intergenic regions.

**Figure S3. VEFS domain of SUZ12 in NAc D1-MSNs does not change levels of H3K27me2 or H3K27me3. (A)** Schematic of experiments involving viral-mediated overexpression of VEFS. **(B)** Representative coronal brain section of a stress-naive mouse showing H3K27me2 expression in GFP-injected (top) or VEFS-injected (bottom) D1-MSNs of the NAc. Scale bars: 20 µm. **(C)**

Percentage of fluorescence intensity for H3K27me2 in GFP-positive and GFP-negative neurons in the NAc. Two-way ANOVA: No significant virus effect: $F_{(1,14)}$=0.004, p=0.95; cell-type effect: $F_{(1,14)}$=0.7, p=0.41 or virus by cell-type interaction: $F_{(1,14)}$=0.71, p=0.42. **(D)** Representative coronal brain section of H3K27me3 expression in GFP-injected (top) or VEFS-injected (bottom) D1-MSNs of the NAc. Scale bars: 20 µm. **(E)** Percentage of fluorescence intensity for H3K27me2 in GFP-positive and GFP-negative neurons in the NAc. Two-way ANOVA: No significant effect of virus: $F_{(1,14)}$=0.43, p=0.52; cell-type: $F_{(1,14)}$=0.086, p=0.77; or interactions: $F_{(1,14)}$=0.087, p=0.77.

**Figure S4. Impaired reversal learning in stress-susceptible. (A)** Schematic of experimental design and timeline. Susceptible (SUS; n=7), resilient (RES; n=6) and control (CON; n=5) groups were trained in the reversal learning task after CSDS. **(B)** No significant differences in the acquisition phase: Two-way ANOVA: group effect: $F_{(2,15)}$=2.026; p=0.16; session effect: $F_{(1.559, 23.39)}$=1.203; p=0.308; group by session interaction: $F_{(4,30)}$=0.579; p=0.68. **(C)** SUS mice exhibit impaired reversal learning: Two-way ANOVA: group effect: $F_{(2,15)}$=16.08; p<0.001; session effect: $F_{(2.864, 42.95)}$= 36.18; p<0.0001, significant group by day interaction: $F_{(15,75)}$=2.342; p<0.01. Tukey's test: Reduced percentage of correct trials in the SUS group as compared to CON and RES, during reversal days 4 (R4), R5 and R6, **p<0.01. **(D)** Positive correlation between the percentage of correct trials during R6 and the interaction ratio. **(E)** Schematic of the *"double-hit"* stress experiment. Mice subjected to ELS further received a second stress exposure by subthreshold social defeat (SbD). **(F)** No differences between the groups during the acquisition phase: Three-way ANOVA: session effect: $F_{(2,21)}$=2.445, p=0.11; ELS effect: $F_{(1,21)}$=0.80, p=0.38; SbD effect: $F_{(1,21)}$=1.63, p=0.21; SbD by ELS by session interaction: $F_{(2,21)}$=0.25, p=0.77. **(G)** Combined exposure to ELS and SbD impaired reversal learning. Three-way ANOVA: session effect: $F_{(5,42)}$=79.67, p<0.0001; ELS effect: $F_{(1,42)}$=9.38, p<0.01; SbD effect: $F_{(1,42)}$=1.53, p=0.22; significant session by ELS interaction: $F_{(5,42)}$=2.46, p<0.05; significant session by SbD interaction: $F_{(5,42)}$=3.19, p<0.05; significant ELS by SbD interaction: $F_{(1,42)}$=14.89, p<0.001. Sidak's comparison for ELS effect: ELS-SbD different from Std-CON at R4, and R6, *p<0.05.

**Figure S5. Chemogenetic inhibition of NAc D1-MSNs impairs behavioral flexibility**. **(A-B)** Schematic of experimental design and timeline. Clozapine N-oxide (CNO) was injected during the reversal training. **(C)** Acquisition: There are no significant differences between groups in the percentage of correct lever presses. Two-way ANOVA: session effect: $F_{(2,51)}$= 4.27; p<0.05; no significant virus effect: $F_{(1,51)}$= 0.034; p=0.85; or interaction: $F_{(2,51)}$=0.179; p=0.83. **(D)** Total number of sessions to reach the learning criterion during the acquisition: $t_{(17)}$=0.90, p=0.37. **(E)** Reversal: Reduced percentage of correct lever presses in mice injected with hM4D(Gi). Two-way ANOVA: virus effect: $F_{(1,102)}$= 7.4; p<0.01, session effect: $F_{(5,102)}$= 15.97, p<0.0001, and no significant interaction: $F_{(5,102)}$=1.25, p=0.29. Sidak's comparison for virus effect: hM4D(Gi) different from mCherry at reversal day 6, *p<0.05. **(F)** Increased number of sessions to reach the learning criterion during the reversal in mice injected with hM4D(Gi): $t_{(17)}$=2.22, p<0.05.

**Figure S6. Overexpression of full-length SUZ12 in NAc D1-MSNs does not change H3K27me1 levels or neuronal intrinsic excitability. (A)** Schematic of the Cre-dependent SUZ12 overexpression construct. **(B)** Schematic of experimental design and timeline. **(C)** Representative coronal brain section of a stress-naive mice showing H3K27me1 expression in GFP-injected (top) or SUZ12-injected (bottom) D1-MSNs of the NAc. Scale bars: 20 µm. **(D)** Percentage of fluorescence intensity for H3K27me1 in GFP-positive and GFP-negative neurons. Two-way ANOVA: No significant effect of virus: $F_{(1,16)}$= 2.69; p=0.12; cell-type: $F_{(1,16)}$= 0.0046;

p=0.94; or interaction: $F_{(1,16)}$=0.0046; p=0.94. **(E)** Current-evoked spikes in D1-MSNs from AAV-GFP- and AAV-SUZ12-injected mice (n= 41-49 neurons/6–8 mice/group) and representative of voltage traces in response to current injections. Two-way ANOVA: current effect: $F_{(30,2640)}$=125.2; p<0.0001. No significant virus effect: $F_{(1,88)}$=0.058; p=0.81; or interaction: $F_{(30,2640)}$=0.253; p>0.99. Sidak's comparison for current effect: ****Different from 100pA, p<0.0001 in GFP and SUZ12. **(F)** Threshold potential: $t_{(88)}$=1.27; p=0.206. **(G)** Resting membrane potential $t_{(88)}$=0.907; p=0.36. **(H)** Percentage of SUS and RES mice in each group. **(I)** Percentage of correct responses during the reversal learning task. Two-way ANOVA: session effect: $F_{(4,70)}$=19.27, p<0.0001. No significant virus effect: $F_{(1,70)}$=0.45, p=0.5; or interaction: $F_{(4,70)}$=0.03, p=0.99. Sidak's comparison for session effect: **Different from R1, p<0.01; $^{\dagger}$Different from R1, p<0.001; in GFP and SUZ12.

Table S1. H3K27me1 enrichment in the SUS vs CON comparison. Related to Figure 2F.

| **GO: Molecular Function** | **Fold Enrichment** | **Enrichment FDR** | **nGenes** |
|---|---|---|---|
| GABA-gated chloride ion channel activity | 9.998 | 0.00006 | 8 |
| Ligand-gated anion channel activity | 6.963 | 0.00017 | 9 |
| Transmitter-gated ion channel activity involved in reg. of postsynaptic membrane | 4.642 | 0.00011 | 14 |
| Cadherin binding | 4.249 | 0.00006 | 17 |
| PDZ domain binding | 3.220 | 0.00011 | 22 |
| Ion channel regulator activity | 2.966 | 0.00019 | 23 |
| Channel regulator activity | 2.910 | 0.00017 | 24 |
| Phosphatidylinositol binding | 2.373 | 0.00009 | 39 |
| GTPase activator activity | 2.036 | 0.00011 | 53 |
| Tubulin binding | 2.003 | 0.00048 | 45 |
| **GO: Cellular Component** | **Fold Enrichment** | **Enrichment FDR** | **nGenes** |
| GABA-ergic synapse | 4.808 | 2.68E-11 | 29 |
| Intrinsic component of postsynaptic membrane | 3.801 | 7.51E-12 | 40 |
| Postsynaptic specialization | 3.106 | 2.81E-18 | 82 |
| Neuron to neuron synapse | 3.097 | 5.09E-18 | 81 |
| Asymmetric synapse | 3.095 | 6.91E-17 | 76 |
| Postsynaptic density | 3.059 | 3.24E-16 | 74 |
| Glutamatergic synapse | 3.051 | 2.59E-19 | 89 |
| Postsynapse | 2.844 | 4.29E-25 | 129 |
| Presynapse | 2.370 | 1.29E-12 | 89 |
| Dendritic tree | 2.315 | 7.00E-15 | 110 |
| **GO: Biological Function** | **Fold Enrichment** | **Enrichment FDR** | **nGenes** |
| Synapse assembly | 3.868 | 9.64E-13 | 45 |
| Synapse organization | 2.960 | 2.19E-16 | 86 |
| Cell junction organization | 2.396 | 1.03E-14 | 109 |
| Trans-synaptic signaling | 2.367 | 1.18E-13 | 103 |
| Synaptic signaling | 2.356 | 5.03E-14 | 107 |
| Neuron projection morphogenesis | 2.353 | 1.62E-12 | 95 |
| Plasma membrane bounded cell projection morphogenesis | 2.352 | 9.85E-13 | 97 |
| Anterograde trans-synaptic signaling | 2.348 | 3.28E-13 | 101 |
| Chemical synaptic transmission | 2.348 | 3.28E-13 | 101 |
| Cell projection morphogenesis | 2.335 | 1.42E-12 | 97 |

Table S2. H3K27me1 depletion in the SUS vs CON comparison. Related to Figure 2I

| GO: Molecular Function | Fold Enrichment | Enrichment FDR | nGenes |
|---|---|---|---|
| Histone methyltransferase activity | 3.259 | 0.0005 | 17 |
| Ubiquitin-like protein-specific protease activity | 2.649 | 0.0003 | 26 |
| Catalytic activity, acting on a tRNA | 2.524 | 0.0004 | 27 |
| Ribonucleoprotein complex binding | 2.291 | 0.0004 | 32 |
| Transcription coactivator activity | 2.040 | 0.0003 | 44 |
| Catalytic activity, acting on a nucleic acid | 2.038 | 0.0000 | 100 |
| Small GTPase binding | 1.962 | 0.0004 | 47 |
| ATP hydrolysis activity | 1.940 | 0.0001 | 59 |
| Microtubule binding | 1.929 | 0.0009 | 44 |
| GTPase binding | 1.916 | 0.0003 | 52 |
| **GO: Cellular Component** | **Fold Enrichment** | **Enrichment FDR** | **nGenes** |
| Membrane coat | 3.095 | 4.83794E-06 | 26 |
| Coated membrane | 3.095 | 4.83794E-06 | 26 |
| Intracellular protein-containing complex | 2.189 | 3.69057E-16 | 138 |
| Glutamatergic synapse | 2.147 | 4.46322E-10 | 90 |
| Golgi membrane | 1.983 | 2.67932E-08 | 88 |
| Endosome membrane | 1.969 | 1.3503E-06 | 70 |
| Transferase complex | 1.969 | 1.41373E-11 | 129 |
| Presynapse | 1.947 | 2.86809E-09 | 105 |
| Nuclear envelope | 1.893 | 1.49767E-06 | 77 |
| Endosome | 1.848 | 7.94086E-11 | 142 |
| **GO: Biological Function** | **Fold Enrichment** | **Enrichment FDR** | **nGenes** |
| Ribosome biogenesis | 3.060 | 2.88E-09 | 48 |
| ncRNA processing | 2.883 | 4.15E-10 | 57 |
| RNA processing | 2.709 | 4.85E-22 | 130 |
| mRNA metabolic proc. | 2.579 | 9.03E-14 | 93 |
| Process utilizing autophagic mechanism | 2.451 | 1.28E-07 | 57 |
| Translation | 2.180 | 6.27E-08 | 76 |
| Modification-dependent macromolecule catabolic proc. | 2.157 | 1.28E-07 | 74 |
| Peptide biosynthetic proc. | 2.142 | 9.34E-08 | 77 |
| Proteolysis involved in cellular protein catabolic proc. | 2.023 | 3.6E-07 | 81 |

Table S3. H3K27me1 enrichment in the ELS-SbD vs Std-CON comparison. Related to Figure 6J

| **GO: Molecular Function** | **Fold Enrichment** | **Enrichment FDR** | **nGenes** |
|---|---|---|---|
| Voltage-gated cation channel activity | 2.657 | 3.49E-05 | 31 |
| Potassium channel activity | 2.607 | 0.000274878 | 26 |
| Transmembrane transporter binding | 2.547 | 7.26E-05 | 31 |
| Cell adhesion molecule binding | 2.131 | 3.49E-05 | 49 |
| Ion channel activity | 2.038 | 9.81E-07 | 72 |
| Passive transmembrane transporter activity | 1.993 | 7.60E-07 | 78 |
| GTPase activator activity | 1.986 | 2.51E-06 | 71 |
| Cytoskeletal protein binding | 1.702 | 2.17E-07 | 142 |
| Inorganic cation transmembrane transporter activity | 1.672 | 0.00020729 | 81 |
| Protein domain specific binding | 1.595 | 7.20E-05 | 106 |
| **GO: Cellular Component** | **Fold Enrichment** | **Enrichment FDR** | **nGenes** |
| Ionotropic glutamate receptor complex | 4.871 | 1.07773E-08 | 21 |
| Neurotransmitter receptor complex | 4.687 | 2.41033E-08 | 21 |
| Integral component of postsynaptic membrane | 3.067 | 1.52437E-09 | 42 |
| Cation channel complex | 2.876 | 8.6339E-11 | 53 |
| Integral component of synaptic membrane | 2.832 | 3.6297E-10 | 51 |
| Postsynaptic density | 2.709 | 1.89507E-16 | 90 |
| Asymmetric synapse | 2.698 | 1.89507E-16 | 91 |
| Neuron to neuron synapse | 2.672 | 1.78278E-16 | 96 |
| Postsynaptic membrane | 2.650 | 1.61015E-12 | 71 |
| Ion channel complex | 2.495 | 6.9332E-10 | 62 |
| **GO: Biological Function** | **Fold Enrichment** | **Enrichment FDR** | **nGenes** |
| Postsynapse organization | 2.957 | 3.89E-10 | 51 |
| Reg. of synapse organization | 2.792 | 1.25E-10 | 59 |
| Synapse organization | 2.707 | 2.38E-18 | 108 |
| Cell morphogenesis involved in neuron differentiation | 2.335 | 1.17E-15 | 119 |
| Neuron projection morphogenesis | 2.308 | 2.27E-16 | 128 |
| Cell junction organization | 2.305 | 2.38E-18 | 144 |
| Plasma membrane bounded cell projection morphogenesis | 2.295 | 2.27E-16 | 130 |
| Axonogenesis | 2.286 | 1.25E-10 | 86 |
| Cell projection morphogenesis | 2.278 | 2.95E-16 | 130 |
